# Magneto-acoustic protein nanostructures for non-invasive imaging of tissue mechanics *in vivo*

**DOI:** 10.1101/2022.05.26.493158

**Authors:** Whee-Soo Kim, Sungjin Min, Su Kyeom Kim, Sunghwi Kang, Hunter Davis, Avinoam Bar-Zion, Dina Malounda, Yu Heun Kim, Soohwan An, Jae-Hyun Lee, Soo Han Bae, Jin Gu Lee, Minsuk Kwak, Seung-Woo Cho, Mikhail G. Shapiro, Jinwoo Cheon

**Author notes:** W.S. Kim and S. Min equally contributed.

## Abstract

Measuring cellular and tissue mechanics inside intact living organisms is essential for interrogating the roles of force in physiological and disease processes, and is a major goal in the field of mechanobiology. However, existing biosensors for 3D tissue mechanics, primarily based on fluorescent emissions and deformable materials, are limited for *in vivo* measurement due to the limited light penetration and poor material stability inside intact, living organisms. While magneto-motive ultrasound (MMUS), which uses superparamagnetic nanoparticles as imaging contrast agents, has emerged as a promising modality for real-time *in vivo* imaging of tissue mechanics, it has poor sensitivity and spatiotemporal resolution. To overcome these limitations, we introduce magneto-gas vesicles (MGVs), a unique class of protein nanostructures based on gas vesicles and magnetic nanoparticles that produces differential ultrasound signals in response to varying mechanical properties of surrounding tissues. These hybrid protein nanostructures significantly improve signal strength and detection sensitivity. Furthermore, MGVs enable non-invasive, long-term, and quantitative measurement of mechanical properties within 3D tissues and organs *in vivo*. We demonstrated the performance of MGV-based mechano-sensors *in vitro*, in fibrosis models of organoids, and *in vivo* in mouse liver fibrosis models.

## Introduction

Tissue mechanical properties are critical regulators of cellular processes driving morphogenesis, tissue homeostasis, and disease progression^1^. Particularly, tissue stiffness is significantly altered during many pathological processes including cancer, diabetes, cardiovascular disease, and fibrosis^2^. The ability to measure the localized tissue stiffness within 3D living, deep tissues is vital to understanding organismal development and disease processes^3^. However, measuring cellular-scale and multi-directional forces within living, three-dimensional tissues remains challenging^4^. A large repertoire of techniques has been developed to measure mechanics at subcellular resolution, but they are limited to *in vitro* measurement of near-surface stiffness in 2D cultured cells or dissected tissues^5–8^. Moreover, measurements at this length scale are different from tissue-level mechanical properties. Macroscopic characterization methods such as traction force microscopy, rheometry, and elastic micropillars have been used to measure forces at supracellular length scale^9–11^.

However, these approaches cannot capture local variations in tissue stiffness, and have only been performed in 2D/3D cell cultures and dissected tissue sections. Shear-wave elastography represents an ultrasound-based method for non-invasive and quantitative assessment of *in vivo* tissue stiffness, but it may have limited resolution making it difficult to detect small tumors, boundary difficulties, and wave speed fluctuation in different tissue types^12^. Recently, several strategies based on injectable and deformable materials such as micrometre-sized oil droplets or hydrogels have enabled quantitative measurement of local mechanical properties within intact soft tissues *in vivo*^4,13–15^. However, these methods rely on fluorescent probes and optical imaging, and thus deploying such biosensors inside deep tissue of living organisms is challenging due to the limited penetration of light in tissue^16^. Furthermore, these materials may be sensitive to local factors, such as pH and temperature, limiting the capability of long-term, *in vivo* measurement of tissue mechanics^17^.

Magnetomotive ultrasound imaging (MMUS) has emerged as a promising ultrasound-based technique capable of measuring mechanical properties of intact tissues *in vivo*^18^. In this method, an external magnetic field induces movement of magnetic nanoparticles (MNPs) and consequently displacement of surrounding tissues, which can be detected by ultrasound imaging, permitting a qualitative inference of mechanical properties of the tissues^19,20^. However, MNPs are sub-optimal contrast agents for ultrasound detection because they have much lower acoustic scattering compared to typical gas-filled ultrasound contrast agents. As a result, MMUS has poor signal-to-noise ratio and low spatial resolution, and requires a high dose of MNPs and bulky magnetic set-up, both of which present challenges in quantification of tissue mechanics *in vivo*^18^.

Gas vesicles are a unique class of air-filled protein nanostructures purified from buoyant photosynthetic microorganisms which act as highly sensitive ultrasound contrast agents for non-invasive imaging due to their unique physical properties and scattering of sound waves^16,21,22^. In this study, we developed magneto-GVs (MGVs) – nanoengineered conjugates of GVs and strongly superparamagnetic nanoparticles. MGVs produce strong MMUS contrast, allowing the direct measurement of local mechanical properties in 3D organoids and living, intact tissues while overcoming the limitations of the current MMUS imaging technique. Based on the principle of MMUS imaging, we hypothesized that we could develop GV-based *in vivo* tissue mechano-sensors that dynamically change their ultrasound contrast in response to differential tissue mechanics. As the mobility of nanoparticles is influenced by the mechanical properties of their surroundings, an increase in surrounding tissue stiffness reduces magnetically induced movement of MGVs and hence decreases MMUS signals. At the same time, the intrinsic ability of GVs to produce nonlinear ultrasound contrast distinct from MMUS provides a measure of MGV concentration in tissues. Therefore, MGV-based MMUS imaging can quantitatively measure various levels of tissue stiffness *in vivo* with enhanced detection limits, sensitivity compared to conventional MMUS contrast agents such as MNPs. With excellent material stability under physiological conditions, MGVs allow for robust and reproducible imaging that is demonstrated in a single application, greatly advancing the *in vivo* capability of MMUS for long-term biological imaging such as monitoring disease progression. In this study, we demonstrated the utility of MGVs for sensitive and quantitative measurement of tissue stiffness through *in vitro* and *in vivo* experiments. We also demonstrated that the unique properties of MGVs could enable long-term disease monitoring and drug screening using *in vitro* 3D organoid and *in vivo* fibrosis models.

## Results

### Development of MGVs and characterization with MMUS imaging

To determine whether the combination of particles with high acoustic contrast and strong superparamagnetism can improve the MMUS imaging capability, we synthesized magnetic nanoparticle-conjugated gas vesicles (MGVs). We established the MMUS imaging system and optimized the system’s magnetic properties using standard magnetic microparticles **(Fig. 1a, Supplementary Fig. 1)**. Based on the MMUS signal intensity, magnetic parameters of 5 Hz, 30 mT, and sine wave were chosen as frequency, max field strength, and temporal pattern of the electromagnetic field, respectively **(Supplementary Fig. 1)**. Zinc-doped iron oxide magnetic nanoparticles (MNPs) were synthesized and functionalized with azide groups, as previously reported^23^. We purified GVs from the cyanobacterium *Anabaena flos-aquae*^24^, and added dibenzocyclooctyne (DBCO) to the GV protein surface through an NH_2_-NHS reaction. MGVs were finally synthesized by conjugating azide-functionalized MNPs and DBCO-GVs via click chemistry **(Fig. 1b)**. After a 4-hr conjugation reaction, MGVs could be isolated from a suspension of unbound MNPs using buoyancy purification **(Fig. 1d)**. MGVs had a hydrodynamic diameter of approximately 494.1±11.5 nm, while functionalized GVs had a diameter of approximately 272±4.5 nm **(Fig. 1c)**. MGVs were stable in different media conditions or during prolonged storage **(Supplementary Fig. 2a-b)**. The conjugation ratio of MNPs to GVs was approximately 186 MNPs per GV **(Fig. 1e, Supplementary Fig. 3)**. The magnetic moment of MGVs was 79 emu/g, which was comparable to that of MNPs **(Fig. 1f)**.

**Figure 1.**
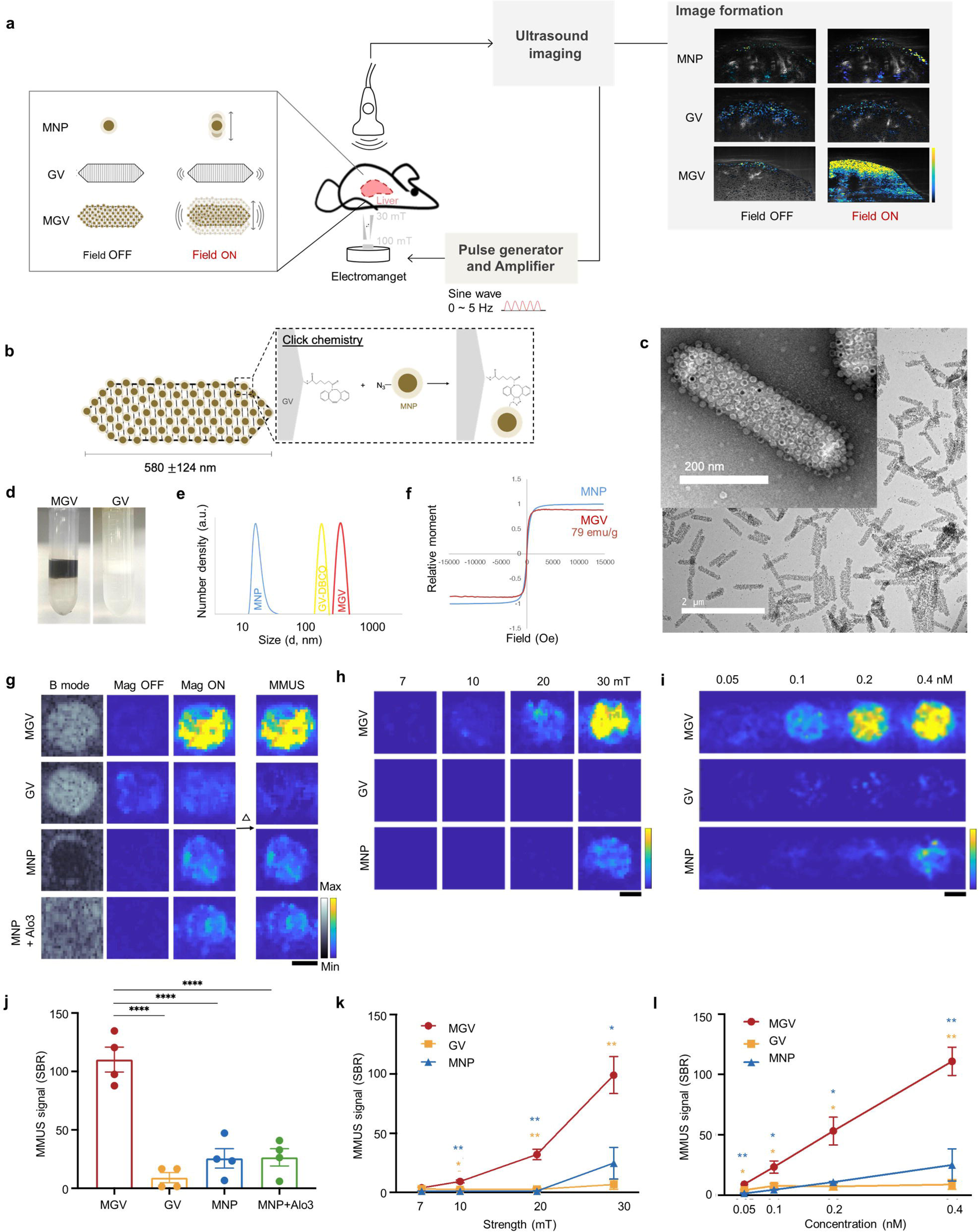
Development and characterization of MMUS imaging system and magnetic nanoparticle-conjugated GVs (magneto-GVs, MGVs). (a) Schematic illustration showing the setup and working principle of the MGV-based MMUS imaging system. (b) Schematic illustration of conjugating magnetic nanoparticles (MNPs) to GVs using click chemistry to form MGVs. (c) TEM image of fabricated MGVs (scale bar = 2 µm, 200 nm). (d) Images of MGVs and GVs after buoyancy purification. (e) The hydrodynamic size of MGVs, MNPs, and GVs with functionalized DBCO by dynamic light scattering (DLS). (f) The magnetic moment of MGVs and MNPs measured by a vibrating-sample magnetometer (VSM). (g) Comparison of MMUS imaging between MGVs, GVs, and MNPs in 0.1% (w/v) agarose phantom using 0.4 nM MGVs, where 0.4 nM MGVs represents 456 pM of GVs and 76 nM of MNPs (*N =* 4 independent samples). (h) Magnetic strength-dependent MMUS images and quantification obtained from 7 mT to 30 mT using 0.4 nM MGVs (*N =* 3 independent samples). (i) Concentration-dependent MMUS images and quantification of different groups (*N =* 3 independent samples). (j-l) Signal to background ratio (SBR) quantification of MMUS images from different cohorts (j), with varying magnetic strength (k) and concentration (l) for the same conditions as in (g-i). For (g-i) scale bars represent 1 mm. Min and max on color bars represent 0 and 10000 arbitrary units, respectively. For (j-l) error bars represent ± SEM, and significance was determined using one-way ANOVA with Tukey’s multiple comparisons test (j) and the multiple unpaired t-test (k, l); *: p<0.05; **: p<0.01; ***: p<0.001; ****: p<0.0001.

We next assessed the ability of MGV protein nanostructures to produce robust MMUS signals in response to applied magnetic fields. Using the optimized magnetic conditions identified above, we performed a head-to-head comparison of GVs, MNPs, and MGVs with MMUS imaging in agarose phantoms. The concentrations of all materials weres matched based on ICP measurements and B-mode imaging **(Fig. 1g, Supplementary Fig. 3)**. MGVs clearly outperformed other nanomaterials in terms of sensitivity and spatiotemporal control, while the spatial resoultion of MMUS imaging largely retained the resolution of conventional B mode ultrasound **(Supplementary Fig. 4)**. Magnetic stimulation of MGVs resulted in robust ultrasound signals, whereas turning off the magnetic field eliminated scattering signals. While both GVs and MGVs produced similar ultrasound contrast using a conventional B-mode sequence, MGVs achieved a 12-fold higher magnetic field-dependent signal intensity change (Δ, 110.1±21.3) than GVs (9.1±8.7) **(Fig. 1g-j)**. In addition, MGVs produced a significant increase in the delta contrast signal compared to MNPs of Fe_2_O_3_ (25.6±16.6) and Al_2_O_3_ (26.5±14.9), suggesting the ability of GVs conjugated to zinc-doped iron oxide MNPs to transduce magnetic stimulation, strongly scatter sound waves, and produce ultrasound contrast for greatly improved MMUS sensitivity **(Fig. 1g-j)**.

To evaluate the performance limits of MGV-based MMUS imaging, we first performed imaging of a dispersion containing MGVs, GVs, and MNPs at a fixed concentration of 0.4 nM in 0.1% agarose while applying magnetic fields with increasing strength ranging from 7 mT to 30 mT. A decrease in magnetic strength was achieved by moving the sample further away from the magnet. MMUS imaging of MGVs revealed a robust non-linear magnetic MMUS response, but not in the negative control of GVs, which produced no visible contrast in response to any level of magnetic field. The MGVs showed a gradual increase of MMUS signal with detectable signals appearing at 10 mT and continuing to increase until 30 mT. In contrast, MNPs produced detectable but substantially weaker MMUS signal only when exposed to a magnetic field ≥ 30 mT **(Fig. 1h-k)**. To assess the sensitivity of the MGV-based approach relative to other nanomaterials, we performed MMUS imaging of a concentration series of MGVs, GVs, and MNPs in agarose phantoms with a constant 30 mT magnetic field. MGVs produced significantly higher delta contrast signals than the other control materials when the concentration was in the range of 0.05 to 0.4 nM **(Fig. 1i-l)**. MGVs attained excellent signal-to-background ratio with a limit of detection (LOD) of 0.05 nM, representing over 8-fold enhancement in sensitivity relative to conventional MNP-based MMUS imaging **(Fig. 1i-l)**. When we compared our results with MGVs to previously published MNP-based MMUS imaging results, we found that MGV-based MMUS uniquely combined sensitivity to relatively low MNP concentrations and relatively weak magnetic fields **(Supplementary Fig. 5)**. Altogether, these results suggest that conjugating GVs with MNPs into a hybrid magnetoacoustic material results in novel contrast agents for MMUS imaging with enhanced ultrasound contrast and detection sensitivity.

### Validation of MGV system in different stiffness conditions

The material stiffness influences the magnetically induced motion of magnetic nanoparticles^19^. Whereas softer materials would allow more MGV movement and induce stronger ultrasound scattering, stiffer materials would restrict MGV motion in response to applied magnetic fields, leading to a decrease in MMUS signal **(Fig. 2a)**. To test the ability of MGVs to quantitatively measure the stiffness of surrounding materials, we performed MMUS imaging *in vitro* of MGVs embedded in agarose phantoms with varying elastic modulus, ranging from 74 Pa to 5828 Pa **(Fig. 2b)**. As predicted, we observed an inverse relationship of MGV signals as a function of increasing elastic modulus of the agarose phantoms. At fixed MGV concentrations (OD_500_ = 4, 0.4 nM) and magnetic field (30 mT), MGVs in 0.1% agarose (109.1±52.3) produced a 2- and 6-fold greater MMUS signal intensity change than in 0.15% agarose (56.9±38.8) and 0.2% agarose (18.7±15.3), respectively, while generating negligible signal in 0.5% agarose phantom (9.3±3.1) **(Fig. 2b-c)**. While the MNP-only sample showed stiffness-dependent signal intensity changes, its MMUS signal was significantly weaker than that of MGVs, leading to lower detection sensitivity of material stiffness **(Fig. 2b-c)**. These results together demonstrate that MGVs embedded in phantoms with lower elastic moduli experience more strain from the same applied magnetic gradient force, resulting in larger vibration amplitudes and stronger ultrasound signals.

**Figure 2.**
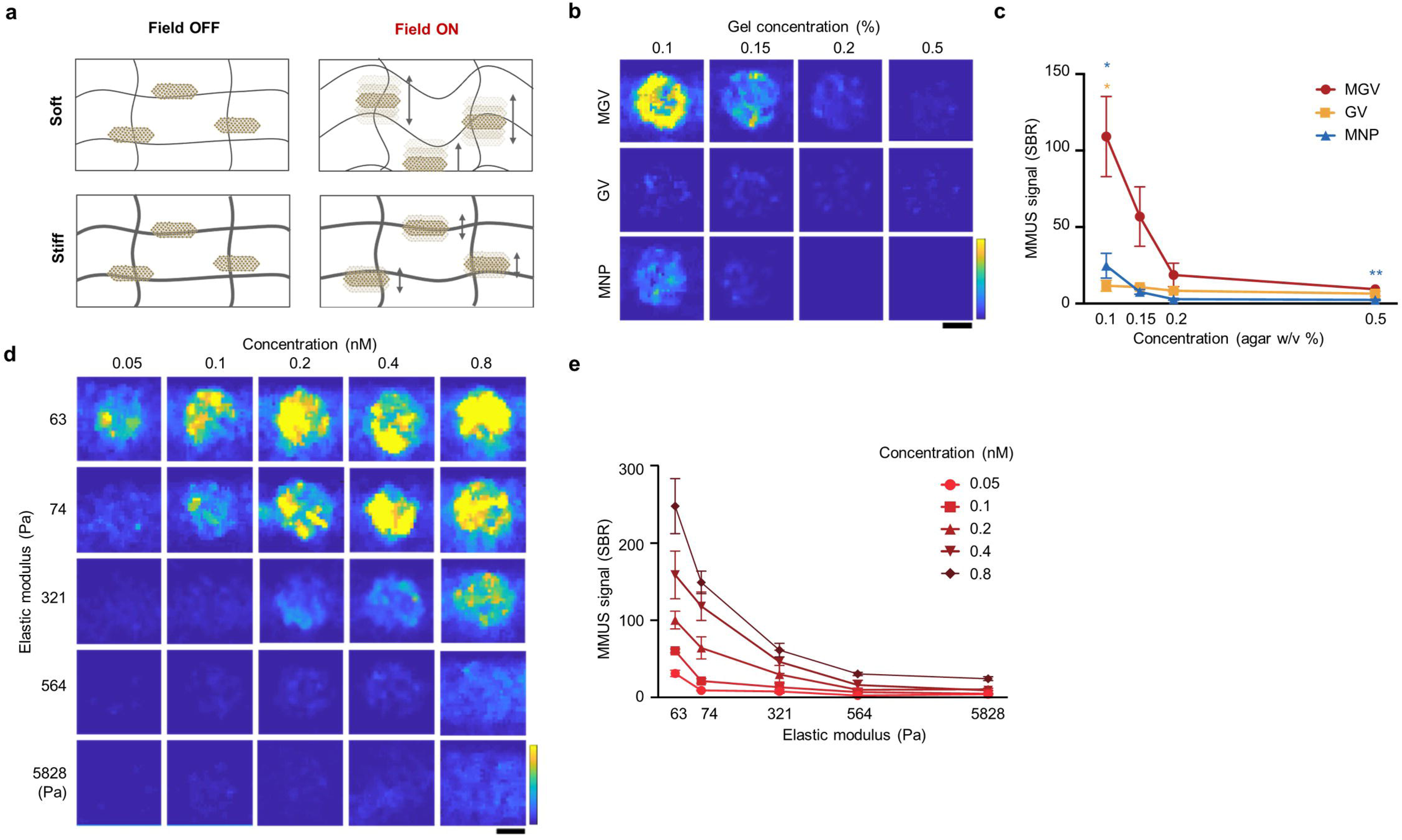
Stiffness-dependent MMUS imaging. (a) Schematic illustration showing the movement of MGVs inside soft and stiff materials when an applied magnetic field is off and on. (b-c) MMUS images and SBR quantification of agarose concentration-dependent movement of MGVs ranging from 0.1% to 0.5% (w/v) agarose (*N =* 4 independent samples). (d-e) MMUS images and SBR quantification based on different values of elastic modulus and concentration (*N =* 5 independent samples). All scale bars represent 1 mm. Min and max on color bars represent 0 and 10000 arbitrary units, respectively. Error bars represent ± SEM, and significance was determined using the multiple unpaired t-test; *: p<0.05; **: p<0.01; ***: p<0.001; ****: p<0.0001.

To determine whether MGVs can more precisely quantify a wide range of material stiffness values, we used two *in vitro* phantoms created from two different hydrogel systems with varying stiffness **(Fig. 2d, Supplementary Fig. 6)**. Matrigel-based phantoms were used for softer materials (63 Pa), and agarose gel phantoms were used for materials with elastic moduli ranging from 74 to 5828 Pa. We observed that the detection range varies with MGV concentration **(Fig. 2d-e)**. The MMUS signal was clearly differentiated in a range of around 63 Pa (100.2±31.7) to 564 Pa (10.0±6.9) when using 0.2 nM MGVs, whereas the detection range broadened to 5,828 Pa (24.3±4.9) when using 0.8 nM MGVs **(Fig. 2d-e)**. Furthermore, when very small amounts of MGVs (0.05 nM) were used, the MMUS signal was only visible until 74 Pa, indicating that at this concentration, MGVs are capable of detecting stiffness changes in materials with elastic moduli less than 74 Pa **(Fig. 2d-e)**. These findings show that depending on the tissue being measured, MGV concentration can be adjusted to generate enhanced or attenuated signals for the acquisition of more accurate MMUS images in diverse tissue types. Moreover, these results demonstrate that MGVs can be utilized as a stiffness sensor, as their signals are connected to the rigidity of their surroundings. In other words, the tissue visualization may vary based on the MGV concentration employed. It would also be feasible to evaluate how the tissue mechanics change over time using the same MGV concentration. To demonstrate this concept, we constructed an organoid model with increasing stiffness in response to the progression of fibrosis.

### Validation of MGV system using lung organoid fibrosis model

Organoids are miniaturized organ-like constructs, and they are in the spotlight as a novel *in vitro* platform to study the development and modeling of disease because they can implement the developmental processes and functional characteristics of actual organs^25^. To confirm the applicability of MGVs in monitoring the tissue- and cellular-level stiffness of 3D microenvironments, a lung fibrosis model was developed with lung organoids. Because lung fibrosis causes tissue stiffening, which leads to a decline in respiratory function and subsequently increased mortality^26^, it is critical to diagnose lung fibrosis at an early stage. Fibrosis was induced by culturing lung organoids with transforming growth factor (TGF)-β to activate TGF-β signaling^27–29^. While normal lung organoids continued to grow in size, fibrotic lung organoids started to decrease in size from day 5 of TGF-β induced fibrosis. Also, we observed that the higher the concentration of TGF-β, the smaller the fibrotic lung organoids became, indicating that the severity of fibrosis was dependent on the level of TGF-β signal activation **(Supplementary Fig. 7)**.

To demonstrate the capability of MGV-based MMUS imaging for sensitive detection of fibrosis progression in a lung organoid model, MGVs were microinjected into the lumen of lung organoids using a Pasteur pipette, and the difference in MMUS signals was compared between normal and fibrosis organoids **(Fig. 3a)**. Microinjection of MGVs conjugated with fluorescence markers showed that MGVs filled the lumen of lung organoids and remained there for 19 days without leakage **(Fig. 3b, Supplementary Fig. 8)**. Consistent with *in vitro* experiments, MGVs produced significantly enhanced MMUS signals compared to GVs and MNPs in the lumen of lung organoids **(Fig. 3c)**. We hypothesized that fibrotic organoids would exhibit an increased stiffness, which would suppress the magnetically induced movements of MGVs and thus result in weaker MMUS signals **(Fig. 3d)**. Lung organoids were moved to a polydimethylsiloxane (PDMS) mold 2 days after MGV microinjection, and the next day fibrosis in MGV-injected organoids was induced with TGF-β treatment using a range of concentrations. MMUS imaging was performed from day 5 to 16 after the induction of fibrosis. The intensity of MMUS signals gradually decreased in the fibrotic lung organoids over a culture period of 5 to 16 days, indicating that lung stiffness increased during the fibrosis progression, and the decrease in signal was more evident in the organoids treated with 50 ng/mL TGF-β **(Fig. 3e-f)**. Our results demonstrate that MMUS imaging using MGVs could provide significant advantages in the detection of fibrosis over other techniques by allowing real-time monitoring of stiffness changes in live lung organoids without fixation.

**Figure 3.**
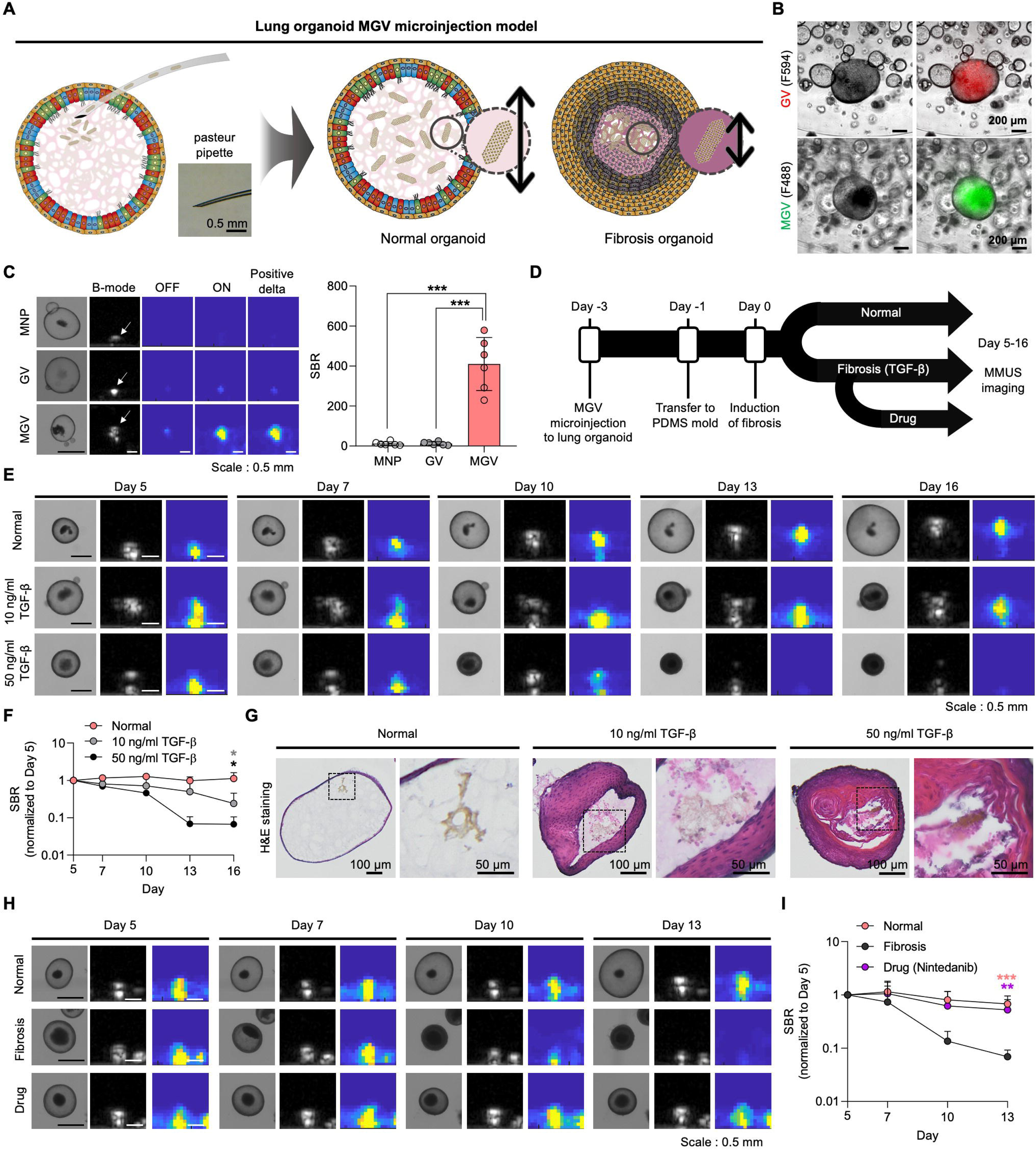
MGV-based MMUS imaging for monitoring fibrosis in human lung organoid model. (a) Schematic illustration showing microinjection of MGVs into a lung organoid and detection of a change in stiffness in a fibrosis lung organoid using MGVs (scale bar = 0.5 mm). (b) Bright-field images and fluorescence merged images of lung organoids microinjected with GVs and MGVs conjugated with Alexa Fluor 594 and Alexa Fluor 488, respectively (scale bars = 200 μm). (c) Bright-field, B mode, and MMUS images of lung organoids microinjected with MNPs, GVs, and MGVs (left panel, scale bars = 0.5 mm) and quantification of the signal to background ratio (SBR) of MMUS images in each group (right panel, *N* = 6). (d) Experimental timeline of the preparation of MGV-microinjected lung organoid models and MMUS imaging. (e) Bright-field, B mode, and MMUS images of MGV-microinjected lung organoids (normal organoid model and fibrosis organoid models treated with 10 or 50 ng/mL TGF-β1). Images were taken from day 5 to day 16 after fibrosis induction (scale bars = 0.5 mm). (f) Quantification of relative SBR signal normalized to the day 5 SBR signal from MMUS images in normal and fibrosis organoid groups (*N* = 3). (g) Hematoxylin & eosin (H&E) staining of organoid sections in normal and fibrosis groups (scale bars = 100 μm and 50 μm in left and right images of each group, respectively). Red arrows indicate the localization of MGVs in the organoids. (h) Bright-field, B mode, and MMUS images of MGV-microinjected lung organoids in the normal group, fibrosis group (50 ng/mL TGF-β1), and drug-treated fibrosis group (50 ng/mL TGF-β1 + 10 μM nintedanib). Images were taken from day 5 to day 13 after fibrosis induction (scale bars = 0.5 mm). (i) Quantification of relative SBR signal normalized to the day 5 SBR signal from MMUS images in normal, fibrosis, and drug-treated fibrosis groups (*N* = 7). All MMUS images in this Figure were obtained using a scale between 0 and 20000 arbitrary units. Error bars represent ± standard deviation (SD), and significance was determined using one-way ANOVA with Tukey’s multiple comparisons test in (c, f, i); *: *p* < 0.05; **: *p* < 0.01; ***: *p* < 0.001.

To validate the increased fibrosis-induced stiffness in lung organoids observed by MMUS, we performed histological and immunohistochemical analyses. Hematoxylin & eosin (H&E) staining revealed thickening of the epithelium layer, abnormal cell growth, and MGV localization in the lumen of TGF-β-treated lung organoids in a dose-dependent manner **(Fig. 3g)**. Moreover, the expression of the known markers that identify ciliated cells (α-tubulin) and goblet cells (MUC5AC) was markedly reduced, while the increased presence of P63-positive basal cells was observed in fibrotic lung organoids with TGF-β treatment **(Supplementary Fig. 9)**. Furthermore, we found a significantly increased expression of smooth muscle actin (SMA) and vimentin (VIM), markers of the epithelial-to-mesenchymal transition (EMT), in basal cells in fibrotic organoids **(Supplementary Fig. 9).** Previous studies showed that in the airway epithelium in patients with idiopathic pulmonary fibrosis (IPF), ciliated cells and goblet cells decrease in number while basal cells proliferate, and the occurrence of EMT in these basal cells results in the formation of fibroblastic foci^30^. These pathological events that form fibroblastic foci in the lumen could lead to increased stiffness in lung organoids. Finally, we examined the feasibility of the MGV-bearing lung organoid model for evaluating the efficacy of anti-fibrosis drugs **(Fig. 3h)**. Nintedanib, well-known for its anti-fibrotic and anti-inflammatory effects on IPF, was tested, and drug treatment was performed starting from the 5^th^ day after fibrosis induction^31^. We found that the intensity of MMUS signals decreased in organoids with TGF-β-induced fibrosis, but the signals in lung fibrosis organoids treated with nintedanib did not decrease and were maintained at a similar level to that of normal organoids **(Fig. 3h-i)**. Based on this result, we verified that MGV imaging in a lung organoid model could be used to screen therapeutic drugs for lung fibrosis.

### MGV-based MMUS imaging in liver organoid fibrosis model

The liver is another important organ for fibrosis modeling due to the high occurrence of hepatic steatosis and cirrhosis. Tissue stiffness is known as an important indicator of the occurrence and progression of fibrotic liver disease^32^. Accordingly, we tested MGV-based MMUS imaging for detecting the increase in stiffness in a liver fibrosis organoid model. As liver organoids do not have a cavity into which MGVs could be injected, an alternative method of MGV incorporation into organoids was considered. Four types of cells (hepatic endodermal cells, hepatic stellate cells, endothelial cells, and mesenchymal cells) were encapsulated in collagen hydrogel containing MGVs, resulting in generation of MGV-incorporated liver organoids **(Fig. 4a)**. Then, fibrosis was induced by TGF-β treatment. Direct encapsulation of MGVs during organoid development allowed localization of MGVs in the extracellular matrix (ECM) in organoids, where the increase in stiffness actually occurs^33^. Because hepatic stellate cells are an important cell type in liver fibrosis and collagen is an ECM component highly correlated with increased stiffness in liver fibrosis, our liver organoid platform contains both cellular and extracellular components suitable for modeling fibrosis. We confirmed the localization of hepatic endodermal cells and hepatic stellate cells in the MGV-incorporated liver organoids, which mimics the cellular compositions of liver tissue **(Fig. 4b)**. Hence, we hypothesized that our liver organoids create a more physiologically accurate model and that MGVs can more accurately sense the increase in stiffness caused by liver fibrosis.

**Figure 4.**
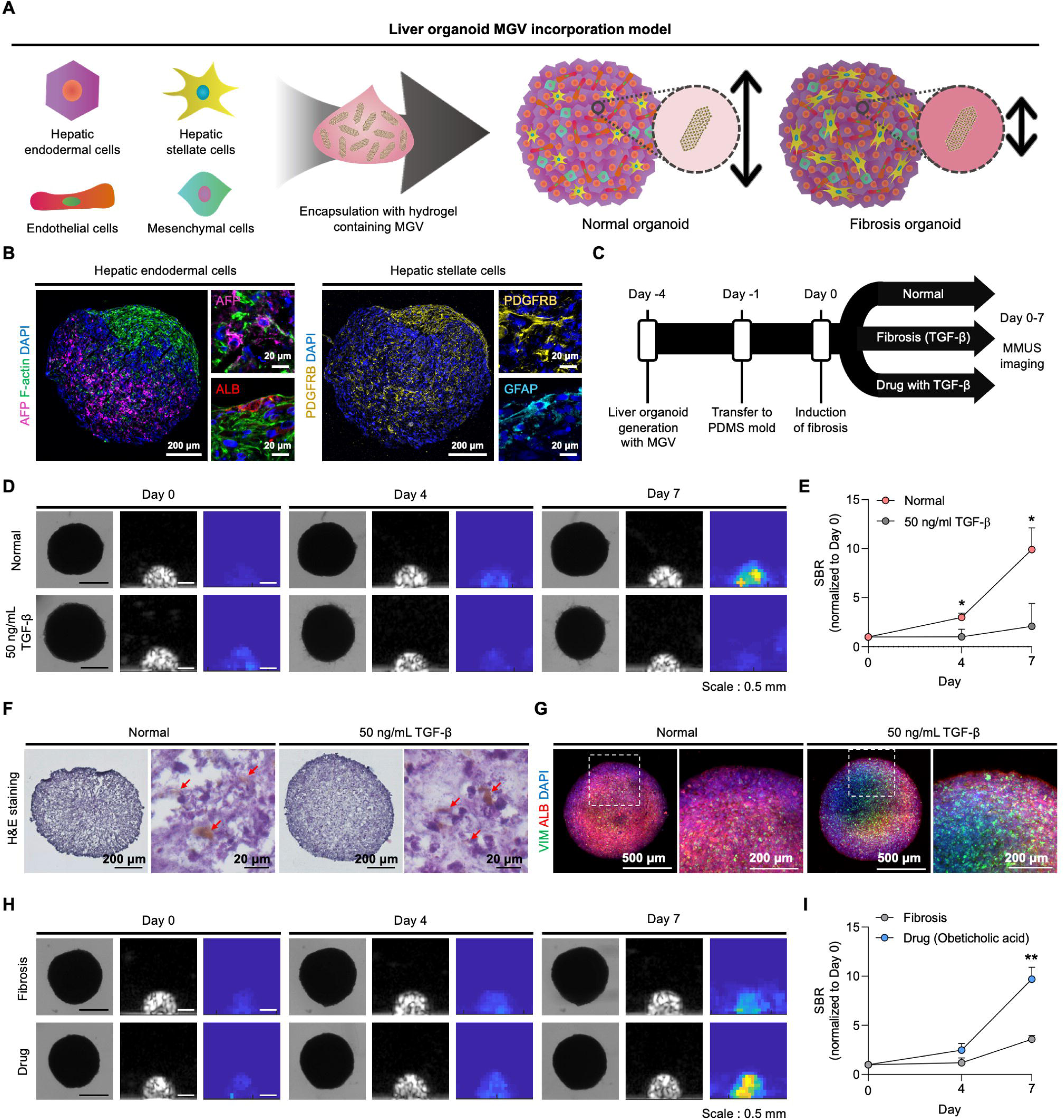
MGV-based MMUS imaging for monitoring fibrosis in human liver organoid models. (a) Schematic illustration showing encapsulation of MGV with four types of cells to generate MGV-incorporated liver organoids and detection of changes in stiffness in fibrosis liver organoids using MGVs. (b) Immunofluorescent images showing the expression of hepatic endodermal cell markers (AFP and ALB) and hepatic stellate cell markers (PDGFRB, GFAP) in liver organoids 3 days after organoid generation. TRITC-conjugated phalloidin was used for cytoskeleton (F-actin) staining and DAPI was used for nuclear staining (scale bars = 200 μm and 20 μm in the left and right images, respectively). The stained signals were presented in pseudo color. (c) Experimental timeline of the preparation of MGV-incorporated liver organoid models and MMUS imaging. (d) Bright-field, B mode, and MMUS images of MGV-incorporated liver organoids (normal organoid model and fibrosis organoid model treated with 50 ng/mL TGF-β1). Images were taken from day 0 to day 7 after fibrosis induction (scale bars = 0.5 mm). (e) Quantification of the relative signal to background ratio (SBR) normalized to the initial signal from MMUS images taken on day 0 in normal and fibrosis organoid groups (*N* = 4). (f) Hematoxylin & eosin (H&E) staining of organoid sections in normal and fibrosis groups (scale bars = 200 μm and 20 μm in the left and right images of each group, respectively). Red arrows indicate the localization of MGVs in the organoids. (g) Immunofluorescent images of fibrotic marker (VIM) and mature hepatic marker (ALB) in normal and fibrosis groups. DAPI was used for nuclear staining (scale bars = 500 μm and 200 μm in the left and right images of each group, respectively). (h) Bright-field, B mode, and MMUS images of MGV-incorporated liver organoids in the fibrosis group (50 ng/mL TGF-β1) and drug-treated fibrosis group (50 ng/mL TGF-β1 + 10 μM obeticholic acid). Images were taken from day 0 to day 7 after fibrosis induction (scale bars = 0.5 mm). (i) Quantification of the relative SBR normalized to the initial signal from MMUS images taken on day 0 in fibrosis and drug-treated fibrosis groups (*N* = 3). All MMUS images in this Figure were obtained using a scale between 0 and 20000 arbitrary units. Error bars represent ± standard deviation (SD), and significance was determined using the unpaired two-sided *t*-test in (e, i); *: *p* < 0.05; **: *p* < 0.01.

Induction of fibrosis in liver organoids was done in the same manner as in the lung organoids, and MMUS imaging was performed from 0 to 7 days after induction **(Fig. 4c)**. Although normal and fibrosis liver organoids did not show morphological differences, there was a significant difference between them in the intensity of the MMUS signals over the culture period **(Fig. 4d-e)**. This result shows that MGVs enable detection of changes in the stiffness of liver tissue. Interestingly, the intensity of the MMUS signal gradually increased in normal liver organoids during culture up to 7 days, indicating that the stiffness of liver organoids naturally decreased over time, which may be attributed to active ECM remodeling during the development of liver organoids^34–36^. In normal tissue, ECM homeostasis is regulated by repeated cycles of ECM degradation and synthesis^34,37^. To verify active ECM remodeling in developing organoids, we checked the expression level of a major collagen component (collagen type I) in liver organoids on days 7 and 10 in organoid culture **(Supplementary Fig. 10a-b)**. Collagen type I content was reduced in 10-day organoids when compared with 7-day organoids. Moreover, considerable expression of matrix metalloproteinase 2 (MMP2) which degrades collagen type I was detected in liver organoids **(Supplementary Fig. 10c)**^38^. Thus, continuous degradation of collagen in liver organoids by MMP2 enzymes secreted from activated cells during development may decrease the stiffness of liver organoids and increase the MMUS intensity in normal liver organoids over the culture period. The presence of MGVs in each organoid model was affirmed through H&E staining of organoid sections **(Fig. 4f)**. The induction of fibrosis was verified through the upregulation of a fibrotic marker (VIM) and the reduction of a mature hepatic marker (albumin, ALB) in liver organoids treated with TGF-β **(Fig. 4g)**. Finally, we examined the applicability of MGV-incorporated liver organoid models to test drugs to treat liver fibrosis. The intensity of MMUS signals in fibrotic liver organoids treated with obeticholic acid, a drug known to prevent or retard liver fibrosis and cirrhosis, was significantly higher than that of the fibrotic organoids without drug treatment **(Fig. 4h-i)**, indicating a significant reduction in stiffness and alleviation of fibrosis by treatment with obeticholic acid. These data demonstrate the possibility of using MGV-incorporated liver organoids as a drug screening platform for liver fibrosis. The combination of MGV-based MMUS imaging and organoid technology could also be utilized for other diseases in which a change in stiffness is an important diagnostic indicator, such as acidosis of the brain and various types of cancers^37,38^.

### MGVs signal detection in animal liver tissues

Having demonstrated the ability of MGVs to serve as MMUS contrast agents and stiffness sensors *in vitro* and *in cellulo,* we next tested their ability to produce robust MMUS signals and measure the stiffness of animal tissues *ex vivo* and *in vivo.* To compare the performance of MGVs, GVs, and MNPs as MMUS contrast agents *in vivo,* we performed intravenous injections of these materials into live, anaesthetized mice. To facilitate more specific GV imaging contrast against tissue background, we modified the protein composition of Ana GVs to generate enhanced non-linear ultrasound contrast under amplitude modulation (AM) pulse sequences^39^. After 5 minutes post-injection, the liver was removed for *ex vivo* MMUS imaging **(Fig. 5a)**. The liver was chosen as our model organ because imaging stiffness would be useful for detecting liver diseases. In addition, GVs naturally accumulate in the liver upon intravenous administration^40,41^. We expected intravenously administered MGVs to be rapidly taken up by the liver, resulting in strong ultrasound contrast in the organ. We observed clear, robust MMUS contrast by MGVs, which exhibited 10-fold stronger signals than those produced by GVs and MNPs **(Fig. 5b-c)**. To demonstrate that MGVs are capable of measuring the mechanical properties of tissues *ex vivo*, MMUS imaging was done in liver samples fixed with 10% formalin for 48 hours^42^. After formalin fixation, the MMUS signals from injected MGVs significantly decreased, corresponding to an increase in tissue stiffness **(Fig. 5b-c)**. Meanwhile, non-magnetomotive AM ultrasound images produced comparable ultrasound conrast from MGVs both before and after formalin fixation, indicating that the decrease in MMUS signal was not due to the collapse or removal of MGVs during fixation, but rather likely due to a restriction of magnetically induced MGV movement **(Supplementary Fig. 11)**. Thus, *ex vivo* liver imaging demonstrated that increasing stiffness lowered the MMUS signal, similarly as in the organoid model.

**Figure 5.**
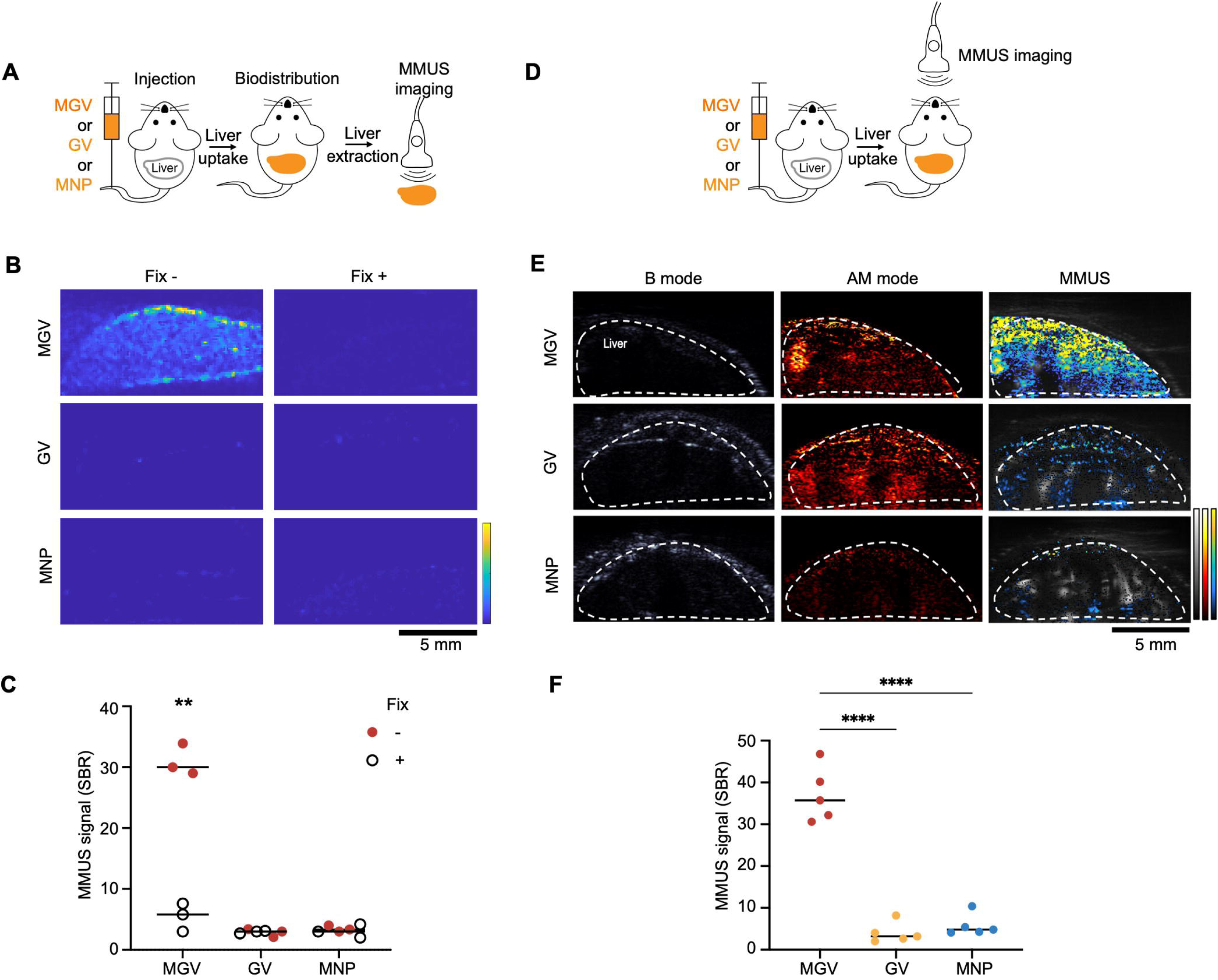
MGV-based MMUS imaging in mouse liver. (a) Experimental scheme of *ex vivo* liver MMUS imaging. (b-c) *Ex vivo* liver MMUS imaging and SBR quantification before and after fixation. MGVs, GVs, or MNPs were injected into the tail vein of mice, and the liver was extracted 5 min after the initial injection. After the first round of imaging, the liver was fixed with 10% formalin for 48 h and imaged again (*N =* 3). (d) Experimental scheme of *in vivo* liver MMUS imaging. (e-f) *In vivo* live animal liver MMUS imaging and SBR quantification. Three different nanomaterials (MGVs, GVs, or MNPs) were injected i.v. and MMUS images were taken after 5 min. B mode images reveal the position of the liver, AM images show the GV signal inside the liver, and Delta MMUS images (parula scale) were overlapped with Doppler images (gray scale) to show the signal below the skin (*N =* 5). All scale bars represent 5 mm. For (b) the color bar represents between 0 and 10000 arbitrary units, and for (e) the min and max on parula and hot color bars represent between 0 and 500000 arbitrary units, respectively, and gray scale bar represents between 0 and 1000000 arbitrary units. Error bars represent ± SEM, and significance was determined using two-tailed heteroscedastic *t*-tests with Welch’s correction within each group (c) and one-way ANOVA with Tukey’s multiple comparisons. *: p<0.05; **: p<0.01; ***: p<0.001; ****: p<0.0001.

Due to ultrasound reflection by the skin and breathing artifacts, which create significant noise and signal attenuation, *in vivo* MMUS imaging of live animals remains challenging. B-mode and Doppler imaging were used to locate the liver, and using ultrafast amplitude modulation (uAM) imaging, we could visualize robust contrast from MGVs and GVs, confirming their delivery to and accumulation in the liver **(Fig. 5e, Supplementary Fig. 12)**. After confirming their localization, we set out to test the ability of MGVs to produce robust ultrasound signals that can be visualized in deep tissues by MMUS imaging to image an *in vivo* biological process within live, breathing animals **(Fig. 5d)**. This is an important challenge in non-invasive, deep tissue imaging of tissue mechanics as it is well-documented that motion artifacts of live animals reduce accuracy and sensitivity^43^. We administered 100 µL MGVs at 2.28 nM into the tail veins of mice, and 5 min later performed MMUS imaging to image their delivery into the liver while animals were being anesthetized. Both GVs and MGVs could be detected with enhanced signals in AM images, while MNPs did not produce any detectable signals *in vivo*. We found that the mean MMUS signal of MGVs (37.1±6.5) was 9.3× and 6.4× stronger than that of GVs (4.0±2.4) and MNPs (5.8±2.6), respectively **(Fig. 5e-f)**. These results demonstrate that MGVs can be used as MMUS contrast agents to improve signal strength and imaging sensitivity in more complex *in vivo* models.

### Detection of liver fibrosis using MGVs

After establishing MGVs as excellent contrast agents for *in vivo* MMUS imaging, we then asked whether our MGV-based system can function as a stiffness sensor to diagnose *in vivo* disease models. We injected C57BL/6 mice with carbon tetrachloride (CCl_4_), a well-established molecule to promote TGF-β signaling and collagen deposition, and induce liver fibrosis^44^. By administering CCl_4_, hepatic stiffness increases concurrently with fibrosis induction, and the stiffness increases until 30 days after injection and then plateaus^45^. After 4 weeks of CCl_4_ treatment, MMUS imaging was performed to assess the mechanical properties of fibrotic and normal livers **(Fig. 6a)**. Intravenously administered MGVs were taken up by liver tissues and retained their ultrasound scattering property, as evidenced by robust ultrasound contrast under AM. While AM imaging confirmed that similar quantities of MGVs were delivered to the livers of both control and fibrotic mice, we observed a significant reduction in MMUS signal in the fibrosis-induced cohort **(Fig. 6b)**. A softness index, which we used as a quantitative indicator of *in vivo* tissue stiffness based on MMUS and AM imaging, was significantly lower in the fibrosis group (4.2±1.0) than in normal controls (14.0±4.6) **(Fig. 6c)**, consistent with previous observations that liver stiffness increases with the progression of fibrosis^45^. Histological and biochemical analyses confirmed the induction of liver fibrosis, as evidenced by pronounced morphological alteration, disruption of tissue architecture, fiber extension, and increased collagen accumulation **(Fig. 6d-f, Supplementary Fig. 13**). No signs of fibrosis or inflammation were observed in control animals injected with MGVs only, suggesting excellent biocompatibility of MGVs **(Fig. 6d-f)**. Our results establish the ability of our magneto-acoustic nanostructures to visualize and quantitatively detect tissue stiffness within the context of *in vivo* imaging. This demonstration suggests that MGVs are useful as contrast agents for the non-invasive, *in vivo* detection of diseases defined by mechanical changes, such as fibrosis and cancer.

**Figure 6.**
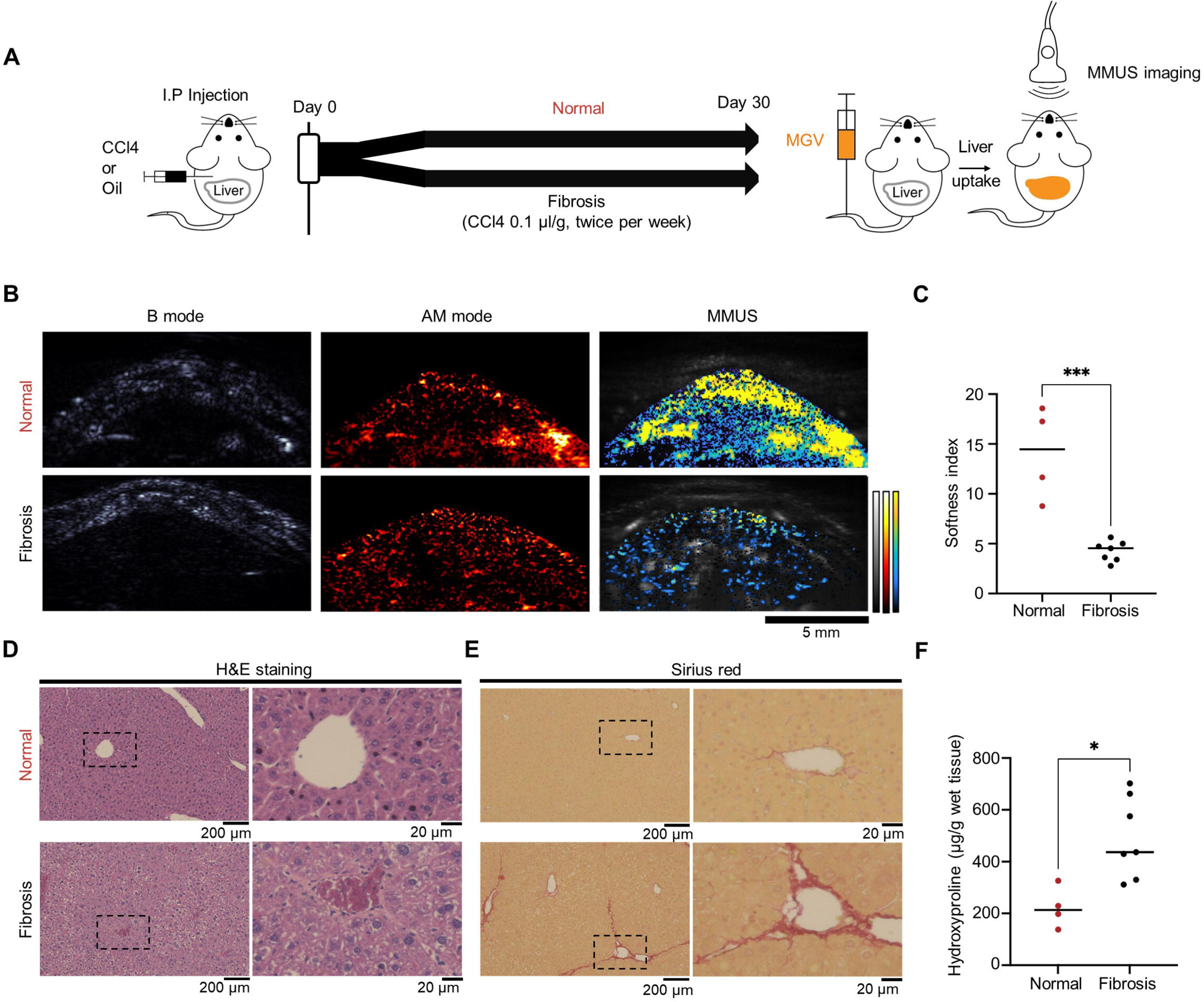
MGV-based MMUS imaging for *in vivo* fibrosis detection. (a) Experimental timeline of inducing liver fibrosis in a mouse model. Either CCl_4_ or mineral oil at a volume of 0.1 µL/g per mouse body weight was injected intraperitoneally twice a week for up to four weeks. (b-c) Ultrasound images and quantification of normal and fibrosis cohorts. The B mode images reveal the position of the liver, AM images show the GV signal inside the liver, and the Delta MMUS images (purula) were overlapped with Doppler images (gray scale) to show the signal below the skin (*N =* 4 for normal group, *N =* 7 for fibrosis group, scale bar = 5 mm). (d) Low-magnification (left) and high-magnification (right) images of H&E stained sections in two groups (scale bars = 200 μm in left images and 20 μm in right images). (e) Low-magnification (left) and high-magnification (right) images of Sirius red-stained sections in two groups (scale bars = 200 μm in left images and 20 μm in right images). (f) Quantification of the hydroxyproline content in liver from two groups. For (b) Min and max in the parula and hot color bars represent 0 and 500000, respectively, and the gray scale bar ranges from 0 to 1000000 arbitrary units. Error bars represent ± SEM, and significance was determined using two-tailed heteroscedastic *t*-tests with Welch’s correction; *: p<0.05; **: p<0.01; ***: p<0.001; ****: p<0.0001.

## Disscussion

Our results establish a new class of hybrid protein nanostructure (MGVs) as nanomaterial-based magneto-acoustically modulated MMUS contrast agents for non-invasive and sensitive imaging and measurement of tissue elasticity *in vivo*. The MGV-based imaging modality couples the improved ultrasound imaging sensitivity of gas vesicles with the capacity of MNPs enabling non-invasive, long-term, safe *in vivo* imaging to deepen our understanding of whole tissue mechanics, compared to prior MMUS imaging or fluorescence-based mechano-sensors^4,18^. The ability of our MGVs to visualize and measure stiffness in 3D tissues *in vivo* was demonstrated by successful detection and monitoring of fibrosis development and therapeutic effects in 3D organoid and *in vivo* models.

Due to rapid uptake and accumulation of nanostructures by the liver, the clinical potential of MGVs for accurate and non-invasive *in vivo* detection of hepatic fibrosis could be readily tested for detecting hepatic fibrosis at an early stage and determining the efficacy of new anti-fibrosis therapies. While a number of non-invasive techniques, such as magnetic resonance imaging (MRI) and shear-wave elastography, have been employed to detect fibrosis, these techniques are also limited by motion artifacts, sampling variability, and insufficient sensitivity^46,47^. Our findings establish magneto-gas vesicles as a promising new class of biosensors for sensitive and non-invasive fibrosis detection *in vivo*. As the elastic modulus of tissues are a reliable biophysical parameter for disease diagnosis^48^, further application-centric optimizations and development of imaging acquisition and processing will help MGV-based imaging achieve widespread use. Additionally, because GV can be genetically encoded inside the cells^49^, combining GV expression with the formation of magnetic cellular nanomaterials, such as magnetotactic bacteria^50^, can home to specific tissues and sense molecular signals in their environment in addition to tissue mechanics. With these improvements, we envision broad utility of our protein nanosturcture as a powerful mechanobiology research platform for quantifying tissue-scale mechanical properties and investigating the roles of these properties during the highly complex processes of development and disease progression.

Recent advances in stem cell-derived organoid systems have provided a promising new class of biological models for understanding various human pathologies and screening of therapeutic drugs and targets^25^. While several recently developed materials could function as *in vivo* force sensors, long-term *in vivo* imaging of 3D mechanics still remains a challenge due to the large size of these materials, their limited stability and low detection sensitivity in deep tissue^4,15^. In this study, MGV-based imaging was successfully used to visualize mechanical dynamics of organoids by monitoring the progression of fibrosis and response to anti-fibrotic drugs for long-term. In particular, the ease of grafting MGVs into organoids by microinjection or encapsulation with the lack of toxicity and degradation, empowers a wide range of mechanobiology studies using organoid models of human diseases. Aa a result, MGV-based measurement of mechanical dynamics may provide novel insights into the roles of mechanobiology in human disease that could be useful in tissue mechanics-based diagnosis and prediction of the best therapeutic outcomes.

## Methods

### Preparation of magnetic nanoparicles (MNPs)

Zinc-doped iron oxide (Zn_0.4_Fe_2.6_O_4_) nanoparticles were synthesized as previously described^23^. Briefly, zinc (II) chloride (ZnCl_2_, ≥98%, Sigma-Aldrich, USA) and an iron (III) acetylacetonate (Fe(acac)_3_, ≥98%, Sigma-Aldrich, USA) were placed in a three-neck round-bottom flask in the presence of oleic acid (Sigma-Aldrich, USA), oleylamine (Sigma-Aldrich, USA), and octyl ether (98%, Aldrich, USA) under argon gas. Silica coating was used to make these nanoparticles water soluble and functionalizable. To begin, the surface was treated with tetraethyl orthosilicate (Sigma-Aldrich, USA), cyclohexane (Deajung, Korea), IGEPAL CO-520 (Sigma-Aldrich, USA), and ammonium hydroxide solution (28%, Sigma-Aldrich, USA) for 24 h at room temperature. The second layer was coated with 3-aminopropyl trimethoxysilane (Sigma-Aldrich, USA) for 2 h. After separation with tetramethyl ammonium hydroxide (TMAOH, 97%, Aldrich, USA), to introduce azide groups on the surface of the nanoparticles, silica-coated nanoparticles (1 mg) were then coated with m-dPEG12-TFP ester (9 mg, Quanta BioDesign, USA), Azido-dPEG12-TFP ester (1 mg, Quanta BioDesign, USA) in DMSO for 2 h at room temperature. Nanoparticles were isolated using a MidiMACS separator column and were dispersed in 10 mM phosphate buffer (PB) buffer.

### Preparation of gas vesicles (GVs)

Anabaena gas vesicles were obtained as previous method^51^. Briefly, GVs were isolated from *Anabaena flos-aquae* using hypertonic lysis and purified using centrifugally assisted flotation. Stripped GVs were prepared by treatment with 6 M urea solution followed by an additional centrifugally assisted flotation and removal of the subnatant. To functionalize DBCO on the surface of the GVs, DBCO-sulfo-NHS ester (Click Chemistry Tool, USA) was mixed with GVs at a molar ratio of 1:10 in deionized water (DIW) for 4 h at 4L at 30 rpm in a vertical shaker. Functionalized GVs were dialyzed in DIW for 72 h with a water exchange every 24 h.

### Development and characterization of MGVs

Magnetic nanoparticle conjugated GVs (magneto-GVs, MGVs) were developed by conjugating MNPs to GVs at a molar ratio of 1:100 for 4 h at 4L at 30 rpm in a vertical shaker. After 4 h, the MGVs were purified three times using buoyancy purification at 300 rpm, 4L, 24 h, and the solvent was replaced with phosphate-buffered saline (PBS) each time. To characterize the morphology, size, and magnetic susceptibility of MGVs, techniques including transmission electron microscopy (TEM, JEOL 2100, JEOL, Japan), dynamic light scattering (DLS, Zetasizer Nano ZS, Malvern, UK), and vibrating sample magnetometer (VSM, Vibration 7407-S, Kake Shore Cryotronics, USA) measurements, respectively, were used. Inductively coupled plasma mass spectrometry (ICPMS, ICAP 7200 Duo + ASX-560, Thermo Fisher Scientific, USA) was used to determine the concentration of MNPs in MGVs. The concentrations of Fe and Zn ions measured by ICP were converted to numbers of MNPs. The concentration of GVs was then calculated by dividing 186 by the average number of MNPs attached to GVs as determined by our TEM images, which was manually calculated **(Supplementary Fig. 3)**.

### Experimental MMUS imaging setup

#### Magnet setup

The schematic illustration of our custom-built MMUS system is illustrated in Fig. 1A. A multipurpose DAQ (USB6003, National instruments, USA), power supply (RSP-1000-24, 24V, 40A, Meanwell), and solid state module (SSR-40DD, FOTEK) were commercially available. To meet the imaging system requirements, the ultrasound system was modified to output a trigger signal prior to imaging to generate amagnetic field. The magnetic field pulse and strength were controlled by using a customized labview system. The field generator was connected to a coil consisting of mutiple turns with a magnetic coil. To increase the magnetic flux density and to localize the magnetic field in the center of the coil, ferritic stainless steel was embedded. The core size was 5 mm in diameter and 100 mm in height. To better focus the field onto a smaller region of interest, a symmetric conic frustum was cut at 56° on the top side. The magnetic pulse strength, measured at 6 mm above the iron-core tip using a digital gaussmeter (DSP 475, Lakeshore Inc., Westerville, OH), was 0.03 T, which was used for the imaging experiments.

#### MMUS imaging and processing

For MMUS imaging, the phantoms were submerged in PBS, and ultrasound images were acquired using a Verasonics Vantage programmable ultrasound scanning system with an L22-14v 128-element linear array transducer with a 0.10-mm pitch, an 8-mm elevation focus, a 1.5-mm elevation aperture, and a center frequency of 18.5 MHz with 67%−6 dB bandwidth (Verasonics, Kirkland, WA). Two sets of ultrasound IQ data were collected for *in vitro* and organoid imaging at each loop containing a pulse sequence consisting of 5 tilted plane waves (varying from −6 to 6 degrees), each containing 500 ensemble coherently compounded frames, collected at a framerate of 500 Hz with a voltage of 3V. A total of 20 loops of images were collected per set. The first set was taken as a background frame for background subtraction with the magnetic field off (Mag OFF). The second set was taken with the magnetic field on (Mag ON), during which the function generator was triggered for 2000 µsec prior to the beginning of the imaging.

To obtain each image, IQ data were processed in the same manner as previously described, with quadrature detection used to extract the generated movement based on the excitation frequency^52^. Briefly, for a set of N frames, let R_I_(x,y,n) + jR_Q_(x,y,n) represent an element in this IQ array with n running from 1 to N, and R_I_ and R_Q_ representing the in-phase and quadrature signal, respectively. First, the received IQ data were phase unwrapped to generate a new 3D array r_unwrapped_(x,y,n) = arg(R_I_(x,y,n) + jR_Q_(x,y,n)). Afterward, quadrature detection was used to tease out the signal that oscillates at the magnetic pulse frequency (f_0_)

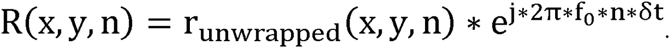

To calculate the displacement amplitude at frequency f_0_ for each pixel, all the frames were averaged to calculate *<u>R</u>*<u>(*x,y*)</u> for the quadrature-detected sequence R(x, y, n). The mean value was used rather than a low-pass filter to determine the displacement amplitude at f_0_, which was obtained as

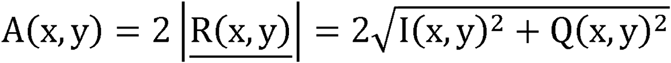

Finally, ultrasound Delta images were constructed by subtracting the Mag ON frame by the Mag OFF frame. For MMUS signal quantification, the Delta images were used. Regions of interest (ROIs) were defined to capture the ultrasound signal from the phantom well or organoid region. All *in vitro* phantom experiments had the same ROI dimensions. For organoid models, ROIs were selected by B mode images in which the organoid size was not same in all cases. Background ROIs were chosen in areas where no sample was present. The mean pixel intensity was calculated for each ROI, and the signal from the background region and sample region was calculated as the signal to background ratio (SBR).

### Rheometer measurement

To prepare the hydrogel samples for the rheometer, agarose or Matrigel was solidified at 25 °C or 37 °C, respectively, at the desired concentration. The rheometer (Anton Paar MCR102) was set to frequency sweep mode from 100 Hz–0.1 Hz (logarithmic, 16 points) and the (ocillatory) shear strain was set to 1%. Elastic modulus values were determined using the storage modulus obtained at 1 Hz.

### Ultrasound phantom preparation

#### *In vitro* phantom

To produce *in vitro* MMUS imaging phantoms, wells were cast with molten 0.5 % w/v agarose in PBS using a custom 3D-printed template. MGV, MNP, or GV (DBCO-functionalized GV) samples were mixed 1:1 with 50 °C agarose and injected into wells prior to solidification. Matrigel was stored at 4°C until loaded and solidified for 30 minutes at 37 °C. Hydrogels and samples were made at a concentration two times greater than the final required concentration.

#### Organoid phantom

For MMUS imaging of organoids, a polydimethylsiloxane (PDMS) mold was fabricated. PDMS solution was prepared by mixing PDMS pre-polymer (Sylgard 184; Dow Corning, Midland, MI, USA) and curing agent (Dow Corning) at a ratio of 10:1 (v/v). Then the mixture was poured into 60 pi petri dishes and cured in a drying oven for 4 h after removing bubbles using a vacuum chamber. The center of the cured PDMS mold was punched to make chambers for the organoids. After sterilizing each PDMS mold with ultraviolet (UV) irradiation for 30 minutes, MGV-microinjected lung organoids or MGV-incorporated liver organoids were encapsulated in growth factor-reduced Matrigel (Corning, Corning, NY, USA) and transferred to the chambers in the mold. After the gelation of Matrigel, the organoids were cultured in growth medium, and MMUS imaging was performed after replacing the medium with 1× phosphate-buffered saline (PBS, Sigma-Aldrich, St. Louis, MO, USA). For fibrosis induction, lung organoids were cultured in medium including recombinant human transforming growth factor (TGF)-β1 (Peprotech, Rocky Hill, NJ, USA) without A83-01 (Tocris, Bristol, United Kingdom). For the drug tests with lung organoids, 10 μM nintedanib (Sigma-Aldrich) was administered to the organoids every 2-3 days starting from day 5 after fibrosis induction. Liver organoids were also cultured in medium containing TGF-β1 for fibrosis induction, and 10 μM obeticholic acid (OCA, Selleck, USA) used for the drug tests was administered to the organoids every 2-3 days starting from the first day of fibrosis induction (day 0).

### Organoid experiments

#### Generation of lung organoids and MGV microinjection

Lung organoids were prepared from human lung tissues harvested with the patients’ consent. The use of human lung tissues for lung organoid generation was approved by the Institutional Review Board (IRB) of Severance Hospital (IRB No: 4-2021-1555). The basal medium used to isolate lung cells from tissues and to prepare growth medium for lung organoid culture consisted of advanced DMEM/F12 (Dulbecco’s Modified Eagle Medium/Ham’s F-12) containing 2 mM GlutaMax, 10 mM HEPES (N-2-hydroxyethylpiperazine-N-2-ethane sulfonic acid) and 1% (v/v) penicillin/streptomycin (P/S) (all from Thermo Fisher Scientific, Waltham, MA, USA). Then, organoid growth medium was prepared using the basal medium supplemented with 10% (v/v) R-spondin1-conditioned medium, 1× B27 (Thermo Fisher Scientific), 100 ng/mL mouse Noggin (Peprotech), 100 ng/mL human fibroblast growth factor-10 (FGF10, Peprotech), 25 ng/mL human fibroblast growth factor-7 (FGF7, Peprotech), 1.25 mM N-acetyl cysteine (Sigma-Aldrich), 5 mM nicotinamide (Sigma-Aldrich), 500 nM A83-01 (Tocris), and 100 µg/mL primocin (InvivoGen, San Diego, CA, USA). R-spondin1-conditioned medium was prepared using HEK293T cells expressing Rspo1-Fc, which were obtained from Calvin Kuo’s laboratory at Stanford University. Lung organoids were generated using a previously reported protocol^53^. Briefly, harvested human lung tissues were fragmented using scissors, and then washed three times with Dulbecco’s phosphate buffered saline (DPBS, Sigma-Aldrich). The tissues were incubated in collagenase type L (2 mg/mL solution in basal medium, Thermo Fisher Scientific) with gentle shaking for 1 h at 37 L. Digested tissues were then washed with DPBS and strongly pipetted with 0.1% (v/v) fetal bovine serum (FBS, Thermo Fisher Scientific) diluted in DPBS to separate cells from the tissue. The suspension was filtered with a 70 μm strainer to remove the tissue debris, and the number of isolated cells was counted. Then, cells were encapsulated in growth factor-reduced Matrigel at a concentration of 5 × 10^6^ cells/mL and were moved into a 48 well plate. After gelation for 10 minutes, growth medium was added to each well. The growth medium was replaced every 3-4 days, and 10 μM Y27632 (BioGems International, Inc., Westlake Village, CA, USA) was added to the culture for the first 4 days. The microinjections of MGVs were performed using a method similar to microinjection into embryos in a previous study^54^. The end of a Pasteur pipette was made narrow for microinjection using an alcohol lamp, and it was connected to a mouthpiece and rubber tubing. MGV solution (OD16) was microinjected into lung organoids, and the organoids were then cultured for 2 days to allow for stabilization. After Matrigel encapsulating the organoids was degraded by treatment with cell recovery solution (Corning), the harvested MGV-microinjected lung organoids were transferred to the organoid chamber of a PDMS mold for MMUS imaging. Microinjection of GVs or MNPs was performed in the same manner as microinjection of MGVs.

#### Human iPSC maintenance

A human induced pluripotent stem cell (hiPSC) line (CHO) was kindly provided by the Yonsei University School of Medicine, and the use of hiPSCs for the liver organoid study was approved by the Institutional Review Board (IRB) of Yonsei University (permit number: 7001988-202104-BR-1174-01E, 7001988-202104-BR-1175-01E). The hiPSC line was characterized for pluripotency and regularly checked for mycoplasma contamination. Culture of hiPSCs was performed using StemMACS iPSC-Brew XF medium (Miltenyi Biotec, Bergisch Gladbach, Germany) under Matrigel-coated, feeder-free conditions. Cells were passaged every 5-7 days using ReLeSR (STEMCELL Technologies, Vancouver, BC, Canada) and maintained within passages 20 to 50.

#### Generation of MGV-incorporated human liver organoids

For generation of liver organoids, four types of cells (hepatic endodermal cells, hepatic stellate cells, endothelial cells, and mesenchymal cells) were prepared. Hepatic endodermal cells and hepatic stellate cells were differentiated from hiPSCs. Human umbilical vein endothelial cells (HUVECs) and human mesenchymal stem cells (hMSCs) were purchased from Lonza (Basel, Switzerland). The differentiation protocol for hepatic endodermal cells was adapted from the previous methods with slight modifications^55,56^. Briefly, hiPSCs were seeded on Matrigel-coated 6-well plates with mTeSR plus medium (STEMCELL Technologies) and 10 μM Y27632. After 24 h, the medium was changed to RPMI 1640 (Thermo Fisher Scientific) and supplemented with 100 ng/mL activin A (R&D Systems, Minneapolis, MN, USA) and 1% B27 minus insulin (Thermo Fisher Scientific), and the cells were cultured for 4 days to induce definitive endodermal cells. CHIR99021 (3 μM, LC laboratory, Woburn, MA, USA) was added to the medium only on the first day of the 4-day differentiation protocol. Then, the cells were further cultured for an additonal 3 days to induce hepatic endodermal cells in RPMI 1640 medium supplemented with 1% B27 (Thermo Fisher Scientific), 20 ng/mL bone morphogenetic protein 4 (BMP4, R&D Systems), and 10 ng/mL fibroblast growth factor-basic (bFGF, Peprotech). The differentiation protocol for hepatic stellate cells was adapted from a previously reported method with slight modifications^57^. Briefly, hiPSCs were seeded on Matrigel-coated 6 well plates with mTeSR Plus medium and 10 μM Y27632. After 24 h, cells were treated with 20 ng/mL BMP4 in a hepatic stellate cell basal medium composed of 40% (v/v) MCDB201 (Sigma-Aldrich), 57% (v/v) DMEM low-glucose (Thermo Fisher Scientific), 1% (v/v) P/S, 0.1LmM β-mercaptoethanol (Sigma-Aldrich), 100 μM L-ascorbic acid (Sigma-Aldrich), 2.5 μM dexamethasone (Sigma-Aldrich), 0.25× insulin-transferrin-selenium supplement (Sigma-Aldrich), and 0.25× linoleic acid (Sigma-Aldrich) for 4 days to induce mesodermal progenitors. Cells were further differentiated into mesothelial cells for an additional 4 days. Cells were treated with 20 ng/mL BMP4, 20 ng/mL fibroblast growth factor-acidic (FGF1, Peprotech), and 20 ng/mL fibroblast growth factor-3 (FGF3, R&D Systems) for the first 2 days and with 20 ng/mL FGF1, 20 ng/mL FGF3, 5 μM retinol (Sigma-Aldrich), and 100 μM palmitic acid (Sigma-Aldrich) for the remaining 2 days. For the final stage of hepatic stellate cell differentiation, 5 μM retinol and 100 μM palmitic acid were added to the hepatic stellate cell basal medium for 4 additional days. The growth medium was exchanged every two days. HUVECs were cultured with EGM-2 medium (Lonza), and hMSCs were maintained with mesenchymal stem cell growth medium (MSCGM, Lonza). To generate MGV-incorporated liver organoids, a 4 mg/mL collagen hydrogel containing MGVs was prepared using rat tail collagen type L (Corning). A pre-gel solution for the collagen hydrogel (final concentration: 4 mg/mL) was prepared by mixing the collagen solution with 10% (v/v) 10× PBS (Sigma-Aldrich) and 10% (v/v) MGVs (OD_500_ = 16). Finally, the pH of the pre-gel solution was adjusted to 7.4 using 0.5 M sodium hydroxide (NaOH, Sigma-Aldrich). After centrifuging a Eppendorf tube containing the four types of cells at a ratio of 10:7:2:2 (hepatic endodermal cell:HUVEC:hMSC:hepatic stellate cell), the supernatant was removed and the pre-gel solution was added for cell resuspension. The final density of total cells was 2×10^7^ cells/mL, and 5 µL of collagen hydrogel was used for the production of one liver organoid. The liver organoids generated in MGV-containing collagen hydrogels were cultured in ultra-low attachment 24-well plates (Corning) using the following medium: basal medium with equal volumes of HCM Bulletkit (Lonza) and EGM-2 medium supplemented with 3% (v/v) FBS, 1% (v/v) P/S, 20 ng/mL HGF (Peprotech), 10 ng/mL Oncostatin-M (ProSpec, Rehovot, Israel), and 0.1 μM dexamethasone.

#### Immunostaining of organoids

Whole-mount immunostaining was performed to check the overall protein expression in organoids. First, cultured organoids were fixed with 10% (v/v) formalin (Sigma-Aldrich) for 1 h at room temperature (RT), and then permeabilized with 0.1% (v/v) Triton X-100 (Sigma-Aldrich) for 30 minutes at RT. After blocking samples with 5% (w/v) bovine serum albumin (BSA, MP biomedical, Asse-Relegem, Belgium) solution for 2 h at RT, the organoid samples were incubated with primary antibodies for 24 h at 4. After washing three times with 1× PBS, secondary antibodies were added to the samples for 24 h at 4. After washing three times with 1× PBS, 4’,6-diamidino-2-phenylindole (DAPI, TCI Chemicals, Tokyo, Japan) was added to stain the nuclei of the organoids. Separately, immunostaining was performed after sectioning the organoids to more clearly identify the internal organoid structure. To this end, cultured organoids were fixed with 10% (v/v) formalin (Sigma-Aldrich) for 1 h at RT, and then optimal cutting temperature (OCT) blocks containing organoids were prepared using OCT compound (CellPath, Newtown, United Kingdom). After slicing the OCT blocks into sections, they were permeabilized with 0.1% (v/v) Triton X-100 for 10 minutes at RT. After blocking the samples with 5% (w/v) BSA solution for 2 h at RT, they were incubated with primary antibodies overnight at 4. After washing three times with 1× PBS, secondary antibodies were added to the samples for 1 h at RT. After washing three times with 1× PBS, DAPI was added to stain the nuclei of the organoid samples. In this study, the following primary antibodies were used: mouse anti-smooth muscle actin (SMA, Santa Cruz Biotechnology, Inc., Dallas, TX, USA), mouse anti-vimentin (VIM, Sigma-Aldrich), rabbit anti-P63 (Abcam, Cambridge, MA, USA), mouse anti-MUC5AC (Abcam), mouse anti-acetylated α-tubulin (α-tubulin, Santa Cruz Biotechnology, Inc.), and rabbit anti-albumin (ALB, Sigma-Aldrich). The following secondary antibodies were used; Alexa-Fluor 488-conjugated anti-mouse IgG (Thermo Fisher Scientific), Alexa-Fluor 594-conjugated anti-mouse IgG (Thermo Fisher Scientific), and Alexa-Fluor 594-conjugated anti-rabbit IgG (Thermo Fisher Scientific). TRITC-conjugated phalloidin (Sigma-Aldrich) for staining F-actin in the cytoskeleton was added to the samples treated with secondary antibodies. Stained organoids and sections were imaged using a confocal microscope (LSM 900, Carl Zeiss, Jena, Germany).

### Animal experiments

All *in vivo* experiments were conducted in accordance with a protocol approved by the California Institute of Technology’s Institutional Animal Care and Use Committee (IACUC). Throughout all injection and imaging procedures, mice were anesthetized with ∼1–2.5 percent isoflurane. Mice were positioned with the liver facing directly upwards. Prior to each experiment, ultrasound gel was centrifuged at 2,000 × *g* for 10Lminutes to remove bubbles, heated to 37L°C, and then carefully applied to the bodies of the mice. To obtain a precise signal, for all *ex vivo* and *in vivo* models, stripped GVs were used. Stripped MGVs were prepared in the same manner as previously described MGVs. The concentration of GVs was matched to the concentration of MGVs. MNP concentration was also matched to the MNP concentration found in the MGVs.

#### *Ex vivo* imaging

For *ex vivo* imaging, three C57 mice aged 8 weeks were injected intravenously in the tail vein with 2,280 pM (OD_500_ = 20) of MGVs and were euthanized after 5 minutes. The concentration used for injections was chosen based on prior research. After euthanasia, the liver was harvested for *ex vivo* imaging. For MMUS imaging, the liver was cast in 0.5% w/v agarose in a 100-mm petri dish and was solidified for 10 minutes. After the first series of imaging, the tissue was fixed for 48 h in 10% formalin at 4. The second series of *ex vivo* imaging occurred after fixation.

#### Live animal imaging

Five 4-week-old C57 mice were intravenously injected with MGVs for *in vivo* imaging. The regions of interest were positioned in the liver tissue using B-mode and Doppler anatomical imaging. The concentration used for injections was chosen based on prior research. MMUS imaging was performed before and after the injection of MGVs with the magnetic field on. 2,280 pM (OD 20) of MGVs were injected intravenously via the tail vein, and MMUS images were taken 5 min post-injection.

#### Fibrosis model

Prior to the *in vivo* fibrosis experiment, animals were randomized between experimental groups; blinding was not necessary. C57 mice were treated with CCl_4_ (1 µL/g body weight, 1:4 dilution with mineral oil, *N =* 7) or with mineral oil alone (1 µL/g body weight, *N =* 4) via intraperitoneal injection two times per week for four weeks^58,59^. After four weeks, the regions of interest were positioned in the liver tissue using B-mode and Doppler imaging. MMUS imaging was performed before and 5 min after injection with the magnetic field on. 2,280 pM of MGVs were injected intravenously via the tail vein in the normal and fibrosis model groups. After MMUS imaging, livers were harvested. Fresh tissue was homogenized and used for a hydroxyproline assay (Sigma-Aldrich, USA). Other parts of the tissue were fixed for 24Lh in 10% formalin and then submerged in 70% EtOH for storage. Next, the fixed tissue was embedded in paraffin, sectioned, and stained with H&E and Sirius red (Abcam, USA).

#### *In vivo* ultrasound imaging

We employed a recently developed method of ultrafast amplitude modulation (uAM) to precisely visualize and quantify ultrasound contrast *in vivo*^40^. Due to the attenuation of applied sound waves caused by the body, we increased the sound pressure to 370 kPa with the same Verasonics system using an L22-14v transducer, which did not collapse either MGVs or GVs ^40^. For each loop, the data were collected at a framerate of 350 Hz with a voltage of 6 V (370 kPa). The pulse sequence consisted of four bursts repeated at three different amplitudes with four different polarity patterns (varying from −14 to 14 degrees). Each burst contained 500 ensemble coherently compounded frames. Two sets of images were taken prior to and following injection. The first set was used as a baseline for background subtraction purposes, with the magnetic field activated prior to injection (before). The second set was taken 5 minutes after injection with the magnetic field activated (after), with the function generator triggered 2000 µsec prior to the start of imaging. A total of twenty looped images were collected per set. We removed frames with poor breathing artifacts based on their Doppler images. To obtain *in vivo* ultrasound images, the same processing procedures were used as previously described, and ultimately, ultrasound Delta images were constructed by subtracting the after image from the before image. For *in vivo* MMUS signal quantification, the Delta images were used. ROIs were selected consistently to exclude edge effects from the skin. Background ROIs was selected where there was no sample at all. The mean pixel intensity was calculated for each ROI, and the signal from the background region and sample region was calculated as the signal to background ratio (SBR). For the fibrosis experiments, the ratio of the MMUS (SBR) signal to the uAM (SBR) signal was calculated and reported as a softness index.

## Statistical analysis

Sample sizes were chosen on the basis of preliminary experiments to have sufficient replicates for statistical comparison. Values and statistical comparisons are given in the text.

## Code availability

MATLAB code is available from the corresponding author upon reasonable request.

## Acknowledgements

This work was supported by the Institute for Basic Science (IBS-R026-D1). This work was also supported by the National Research Foundation of Korea (NRF) grant funded by the Korea government, the Ministry of Science and ICT (MSIT) (No. 2021R1A2C3004262) and Samsung Research Funding & Incubation Center of Samsung Electronics under Project Number SRFC-TC2003-03. M.G.S. is an Investigator of the Howard Hughes Medical Institute (HHMI).

This article is subject to HHMI’s Open Access to Publications policy. HHMI Investigators have previously granted a nonexclusive CC BY 4.0 license to the public and a sublicensable license to HHMI in their research articles. Pursuant to those licenses, the author-accepted manuscript of this article can be made freely available under a CC BY 4.0 license immediately upon publication.

## Contributions

W.-S.K. performed overall *in vitro* and *in vivo* experiments and analyzed data. S.M designed and performed organoid related experiments. S.K.K., Y.H.K., and S.A. supported hiPSC differentiation, MMUS imaging, and hydrogel characterization, respectively. S.K. provided magnetic nanoparticles. H.D. and A.B. performed initial setup and discussions on this research. D.M. provided gas vesicles. J.-H.L. helped initial setup and discussions on this research. S.H.B. and J.G.L. provided lung tissue for the organoid development. M.K. assisted with the design of research and experiment. W.-S.K and M.K. wrote the manuscript with input from all authors. S.-W.C., M.G.S., and J.C. conceived and supervised the project.

