## Supplementary figures for "Magneto-acoustic protein nanostructures for non-invasive imaging of tissue mechanics *in vivo*"

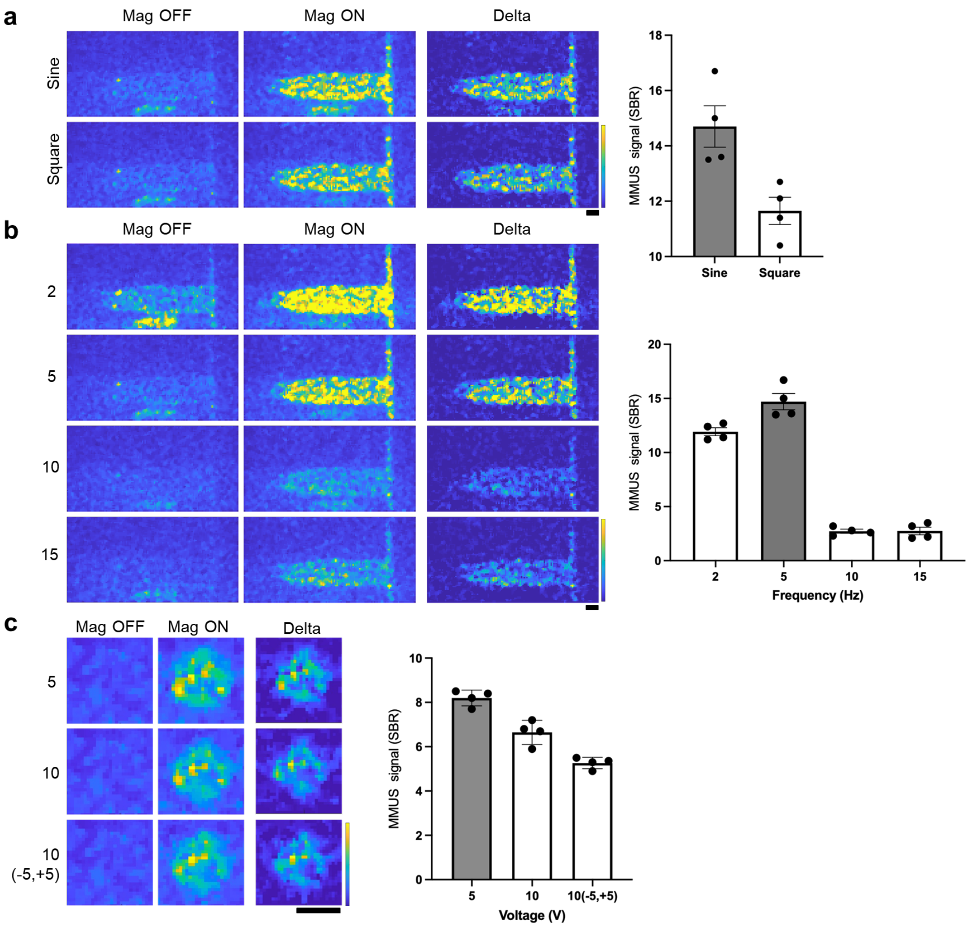


**Supplementary Figure 1. Optimization of magnetic properties using 5-micrometer magnetic particles.** (a) MMUS images and quantification of the sine and square wave forms. (b) MMUS images and quantification using different magnetic frequencies. (c) Images and quantification using different magnetic amplitudes. All scale bars represent 1 mm. Min and max on the parula color bar represent 0 and 2000 arbitrary units, respectively. Error bars represent ± SEM.

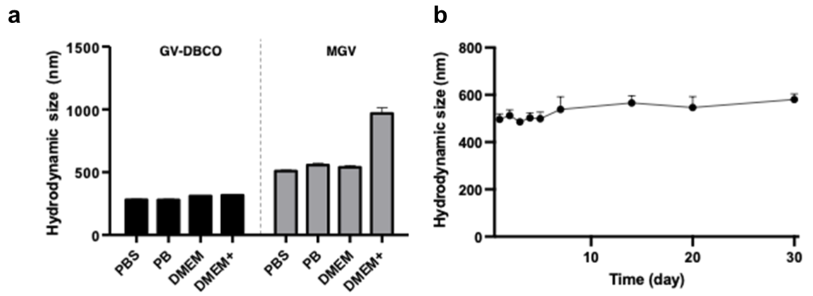


**Supplementary Figure 2. Stability of MGVs in various media.** Dynamic light scattering (DLS) analysis indicating the hydrodynamic size of MGVs (a) in different media conditions incubated for 1 h at 37°C and (b) in aqueous solution for 30 days at 4°C (*N* = 18, number of trials = 3). Error bars represent ± SEM.

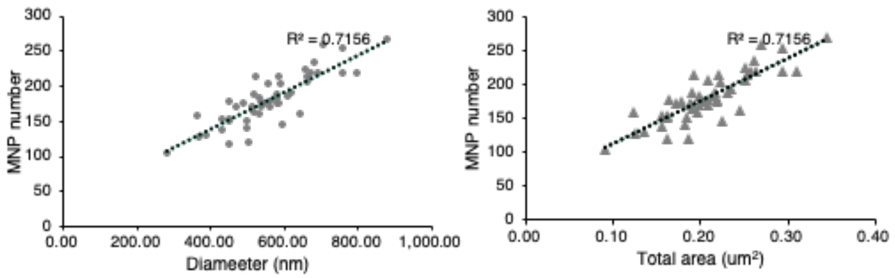


| | MNP Number | GV |
|---|---|---|
| Diameter | 186±34 | 581.3±124.7 nm |
| Area | 870 ± 116 | 1 μm³ |

**Supplementary Figure 3. Quantifying MNP numbers in MGVs.** Average MNP numbers in MGVs per GV diameter or area were calculated based on TEM images. MNPs were manually counted in individual TEM images of MGVs. ImageJ was used to calculate the GV diameter (*N* = 50).

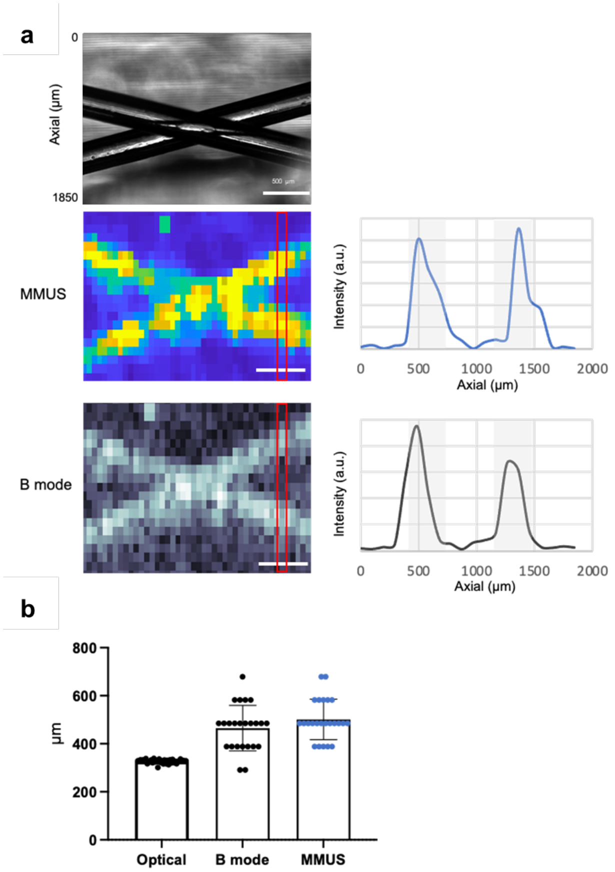


**Supplementary Figure 4. Spatial resolution estimation of MMUS and B mode imaging using an agar phantom with two cross lines of MGVs.** (a) Representative bright-field optical imaging of an agar phantom. The ground truth of the phantom thickness was quantified using ImageJ. Phantom MMUS and B mode images, and the intensity profiles, are shown for a vertical orientation. Scale bar: 500 µm. (b) Average line thickness of optical, B mode, and MMUS images ($N = 24$). Error bars represent ± SD.

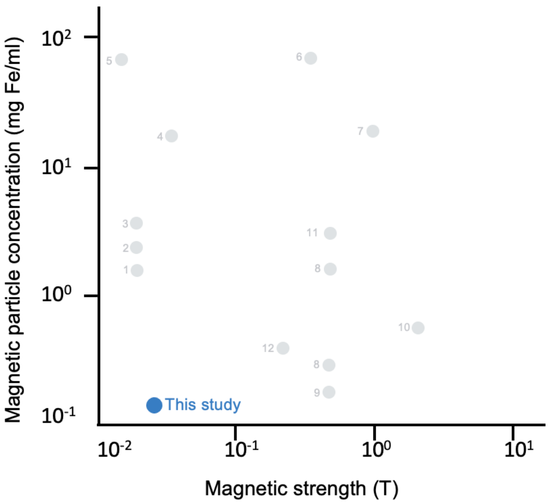


**Supplementary Figure 5. Comparison among previous MMUS studies.** The minimum concentration of magnetic particles (mg Fe/mL) and the minimum magnetic strength to achieve MMUS imaging in previous methods were compared to this study[1–12].

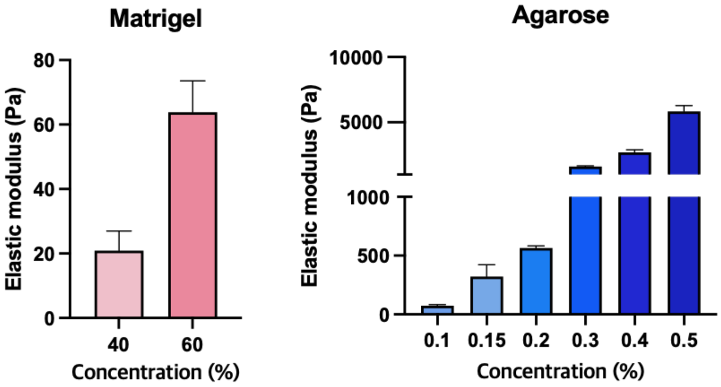


**Supplementary Figure 6. Elastic modulus of hydrogel materials.** Two types of hydrogel materials, Matrigel and agarose, were solidified using desired concentrations at 37 °C and 25 °C, respectively. The elastic modulus were determined using the rheometer storage modulus obtained at 1 Hz.

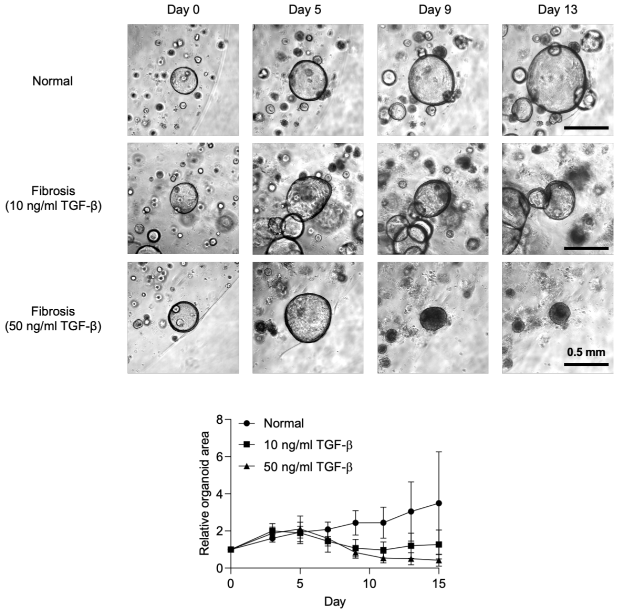


**Supplementary Figure 7. Tracking the fibrotic response of lung organoids during culture.** Bright-field images and size quantification graph of lung organoids composed of one normal model and two fibrosis models treated with 10 or 50 ng/mL TGF-β1. Images were taken from day 0 to day 15 after fibrosis induction (scale bars = 0.5 mm). Error bars represent ± SD.

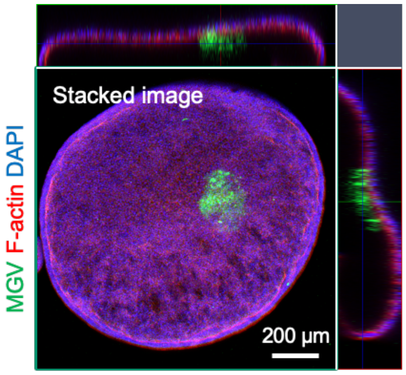


**Supplementary Figure 8. Location of MGV microinjected to lung organoids.** Stacked fluorescent image of a lung organoid microinjected with Alexa Fluor 488-conjugated MGVs 19 days after microinjection. F-actin was used for cytoskeleton staining, and DAPI was used for staining nuclei (scale bar = 200 μm).

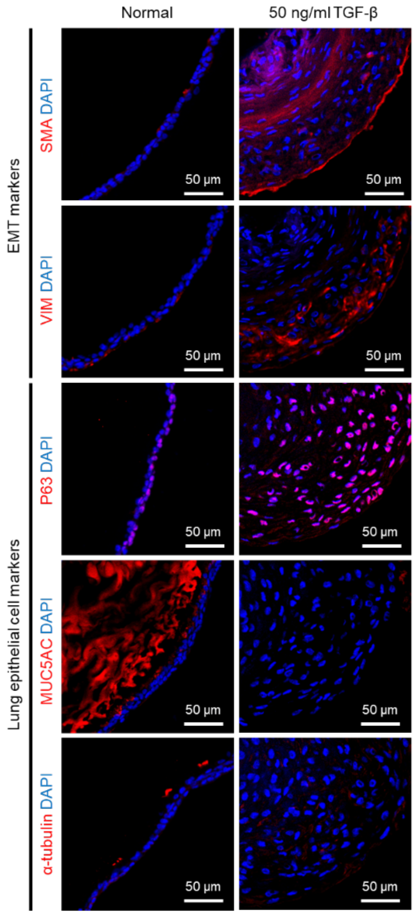


**Supplementary Figure 9. The comparison of protein expression between lung normal organoids and lung fibrosis organoids.** Fluorescent images of MGV-microinjected lung organoids stained with EMT markers (VIM and SMA) and lung epithelial cell markers (α-tubulin, MUC5AC, and P63) 16 days after fibrosis induction by treatment with 50 ng/mL TGF-β1. DAPI was used for staining nuclei (scale bars = 50 µm).

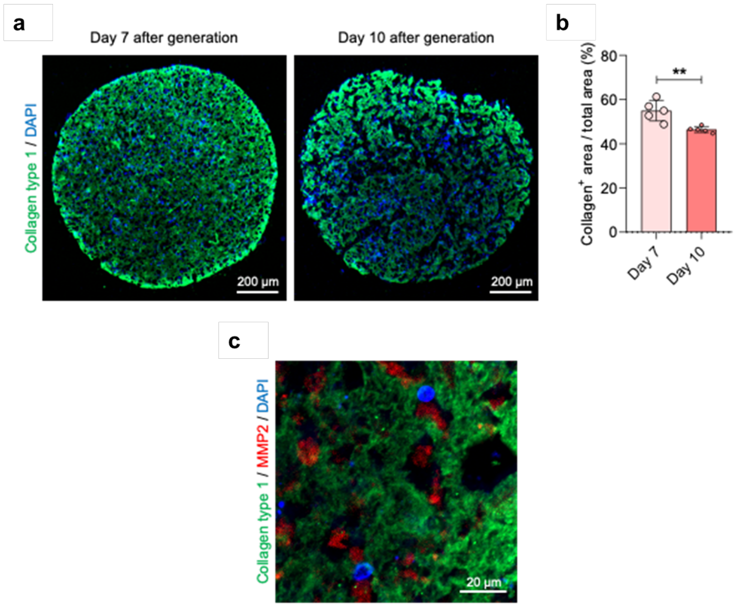


**Supplementary Figure 10. ECM remodeling in liver organoids.** (a) Fluorescent images of normal liver organoid sections stained with collagen type 1 on days 7 and 10 after organoid generation. DAPI was used for staining nuclei (scale bars = 200 μm). (b) Bar graph quantifying the collagen$^+$ area / total area (%) of liver organoids on days 7 and 10 after organoid generation ($N$ = 5). Error bars represent ± SD, and significance was determined using the unpaired two-sided t-test; **: $p<0.01$. (c) Fluorescent image of normal liver organoid sections stained with collagen type 1 and MMP2 on day 7 after organoid generation. DAPI was used for staining nuclei (scale bar = 20 μm).

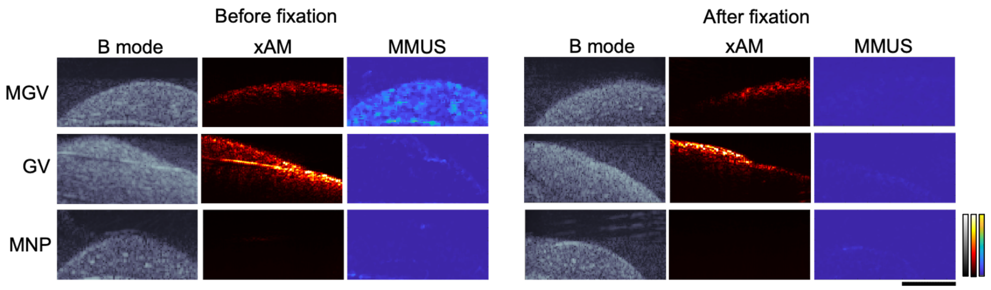


**Supplementary Figure 11. *Ex vivo* liver imaging before and after fixation.** Representative images of *ex vivo* liver imaging. Three different groups of nanomaterials (MGVs, GVs, or MNPs) were injected intravenously into different cohorts, and after 5 minutes, the liver was extracted to take ultrasound images. Liver was fixed in 10% formalin for 48 h. B mode images reveal the shape of the liver, xAM images show the GV signal located at the ultrasound focus, and the Delta images show the MMUS signal. Scale bars represent 5 mm. The scale bars for parula and gray scale range from 0 to 10000 arbitrary units, and the hot scale bar ranges from 0 to 1000000 arbitrary units.

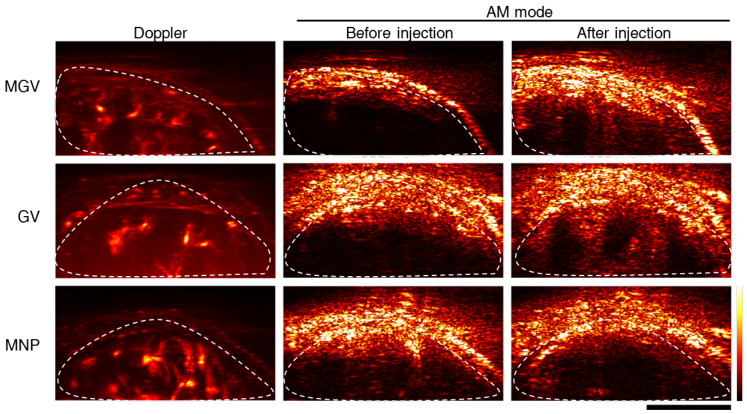


**Supplementary Figure 12. Tracking GV and MGV delivery using AM and Doppler imaging in the liver.** Three different groups of nanomaterials (MGVs, GVs, or MNPs) were injected intravenously into different cohorts, and liver AM and Doppler images were taken after 5 minutes. Doppler images reveal the position of the liver, AM images show the signal from GVs or MGVs after injection. Scale bars represent 5 mm. The color bar ranges from 0 to 1000000 arbitrary units.

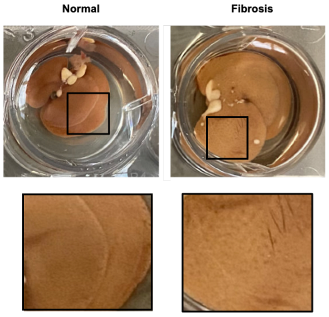


**Supplementary Figure 13. Differences in liver morphology between normal and fibrosis-induced liver organoid models.** Representative image of the liver for normal and fibrosis models treated with either mineral oil or $CCl_4$ (1 μL/g body weight), respectively. The liver surface was punctured and scarred on the outer surface of the liver organoid in the fibrosis group.
